# DLRNA-BERTa: A transformer approach for RNA-drug binding affinity prediction

**DOI:** 10.1101/2025.09.05.674445

**Authors:** Pasquale Lobascio, Khalid Saeed, Asifullah Khan, Ziaurrehman Tanoli

**Affiliations:** Institute for Molecular Medicine Finland (FIMM), HiLIFE, University of Helsinki, Finland; King Abdullah International Medical Research Center (KAIMRC), King Saud bin Abdulaziz University for Health Sciences; Pakistan Institute of Engineering & Applied Sciences (PIEAS), Islamabad, Pakistan; BioICAWtech, Helsinki, Finland

## Abstract

RNA-based therapies are a rapidly expanding field, offering treatments for a wide range of diseases, including many rare conditions. To date, 24 RNA therapeutics have received FDA approval, with 131 more in clinical trials, underscoring RNA’s growing role in modern medicine. In this context, Bidirectional Encoder Representations from Transformers (BERT) models provide a cost-effective and accurate virtual screening strategy for accelerating RNA-targeted drug discovery. These models take RNA FASTA sequences and compound SMILES strings as inputs and generate predicted binding affinities in nanomolar units.

In this study, we introduce DLRNA-BERTa, a RoBERTa-based framework combining RNA-BERTa, pretrained on 9.76 million RNA sequences, with ChemBERTa-v2 for predicting small molecule–RNA interactions. The framework includes six class-specific models, aptamers, repeats, ribosomal RNAs, riboswitches, microRNAs (miRNAs), and viral RNAs, plus a general model for cases where the RNA class is unknown. Proposed DLRNA-BERTa consistently outperforms existing RNA–drug interaction prediction methods. Pearson correlation coefficients achieved are: 0.94 (aptamers), 0.95 (repeats), 0.93 (ribosomal RNAs), 0.94 (riboswitches), 0.95 (viral RNAs), 0.98 (miRNAs), and 0.92 (general model), demonstrating robust performance across RNA classes. Benchmarking against four independent datasets from the ROBIN repository further confirms generalizability.

Application of DLRNA-BERTa to 3,492 approved drugs from the ChEMBL database identified 2,859 compounds with predicted affinities (pKd ≥ 6) across 294 RNA targets. As proof of concept, bleomycin is highlighted, supported by literature evidence of RNA-binding activity. A publicly accessible web application is available at https://huggingface.co/spaces/IlPakoZ/DLRNA-BERTa, in alignment with FAIR principles.

## 1. INTRODUCTION

Over the past three decades, RNA therapeutics are established as a transformative modality in modern medicine, enabling precise modulation of gene expression for the treatment of rare genetic disorders and the development of advanced vaccines, such as those against SARS-CoV-2 (COVID-19) [1,2]. Because of their molecular specificity, RNA-based drugs are considered to provide significant advantages over traditional small molecules and protein therapeutics. These advantages are reflected in the ability to silence disease-associated genes through antisense oligonucleotides or RNA interference (RNAi), as well as to promote therapeutic protein expression through the delivery of synthetic messenger RNA (mRNA) [3,4]. In this way, RNA therapeutics are recognized as capable of addressing a wide range of intracellular and nucleic acid–associated targets, including many previously regarded as “undruggable” [5].

The increasing maturity of this therapeutic class is reflected in regulatory approvals. By 2023, 22 RNA-based drugs are approved by the FDA and EMA [6], and two further approvals are reported in 2024. Olezarsen is approved as an antisense oligonucleotide targeting *APOC3* mRNA to suppress apolipoprotein C-III expression for the treatment of familial chylomicronemia syndrome (FCS) [7]. In parallel, imetelstat is approved as a lipid-conjugated oligonucleotide that binds the telomerase RNA component (*TERC*), providing therapeutic benefit in lower-risk myelodysplastic syndromes (MDS) with transfusion-dependent anemia [8]. By inhibiting telomerase activity, the replicative potential of malignant hematopoietic cells is limited, underscoring the ability of RNA therapeutics to modulate non-coding RNA targets.

At the same time, advances in artificial intelligence (AI) and computational methods are revolutionizing biomedical data analysis. With the increasing availability of high-throughput datasets in genomics, transcriptomics, and drug screening, AI is positioned as a key driver in drug discovery pipelines [9]. Applications are reported in drug repurposing [10,11] and de novo drug design [12,13], leading to more efficient discovery and reduced development timelines.

Drug–target interaction (DTI) is regarded as a central task in drug discovery, aimed at estimating binding affinity between drug-like molecules and biological targets. Accurate DTI models are enabling high-throughput virtual screening, thereby facilitating novel compound identification and drug repurposing. For instance, Aron et al. [14] employed an attention-based model to screen hundreds of thousands of compounds across 1,251 human proteins. Similarly, other transformer-based methods like MolTrans [15] and DLM-DTI [16] frame DTI prediction as a classification task, leveraging sequence embeddings to capture complex drug–target relationships. Despite these advances, RNA targets are receiving comparatively less attention in DTI prediction. This gap is attributed to the scarcity of high-quality RNA–ligand datasets and the historic prioritization of protein targets [17]. Nevertheless, recent efforts are beginning to bridge this gap. DeepRSMA [18] is introduced as a cross-fusion transformer to integrate RNA and compound features, while Xia et al.[19] describe a geometric deep learning approach applied to 3D RNA–ligand complexes, achieving a pearson correlation of 0.67. A review by Zhuo et al. [20] summarizes progress in this domain and highlights methodological directions. Among early benchmarks, a regression-based model proposed by Krishnan et al. [21] represents an important benchmark. Their approach involves a multivariate regression model built from engineered features of RNA targets and small molecules. Despite its simplicity, the model achieves a mean pearson correlation of ∼0.83 on the validation set. Additionally, by applying a binding affinity threshold (pKd ≥ 4.0), they report an F1-score of ∼0.97 on an external classification dataset, demonstrating strong generalization. A related study by Bae et al. [22], currently under review, introduces a sequence-based DL model that jointly predicts RNA–drug interactions and RNA binding sites, reflecting a growing research interest in this area.

In this work, DLRNA-BERTa is introduced as a Large Language Model (LLM) designed to predict RNA–drug binding affinities from primary molecular representations: RNA nucleotide sequences and compound SMILES strings. Our architecture integrates ChemBERTa-v2 [23] with RNA-BERTa, a transformer pretrained on 9.76 million RNA sequences using masked language modeling. RNA and compound embeddings are combined through a cross-attention mechanism, and binding affinities are predicted using a linear output layer. Inspired by DLM-DTI [16], enhancements such as cross-attention and μParametrization (μP)[24] are adapted. By contrast, Low-Rank Adaptation (LoRA) [25] and LoRA-Fine-Tuning-aware Quantization (LoFTQ) [26] do not improve efficiency and are excluded. DLRNA-BERTa is shown to outperform the benchmark model of Krishnan et al. [21], achieving superior performance across six RNA classes and the general model. The trained model is subsequently applied to virtually screen 3,492 FDA-approved drugs, leading to the identification of multiple approved drugs predicted to bind RNA targets. Notably, bleomycin is highlighted through literature validation as interacting with microRNAs and repeat RNA elements, confirming both the biological relevance and predictive reliability of the framework. To support accessibility, a graphical user interface (GUI) and an API are developed, available at https://huggingface.co/spaces/IlPakoZ/DLRNA-BERTa, and source code and datasets are openly available at https://github.com/IlPakoZ/rnaberta-dti-prediction, in accordance with FAIR principles.

## 2. MATERIAL AND METHODS

### 2.1. Pre-training datasets

The pretraining dataset comprises 10,841,246 RNA sequences, partitioned into 9,757,119 sequences for training and 1,084,127 sequences for validation. Collectively, these sequences represent approximately 1.22 billion tokens, of which ∼1.07 billion are assigned to the training set. RNA sequences are obtained from RNAcentral [27] and NCBI [28], with individual sequence lengths capped at 2,000 nucleotides. To ensure consistency with downstream applications, only RNA classes present in the fine-tuning set, aptamers, repeats, ribosomal RNAs, riboswitches, ribozymes, and viral RNAs, are retained. The class-level distribution of the pretraining dataset is as follows:

- 7,979,027 ribosomal RNA (rRNA) sequences, extracted from the SILVA database [29]
- 492,955 microRNA (miRNA) sequences (including both pre-miRNA and mature forms), extracted from RNACentral database
- 3,137 repeat RNA sequences, extracted from RNACentral database
- 90,467 riboswitch sequences, extracted from RNACentral database
- 29,581 ribozyme sequences, extracted from RNACentral database
- 2,246,079 viral RNA sequences, extracted from the NCBI virus database

The mean sequence length across the dataset is 901 nucleotides. To reduce the risk of data leakage between pretraining and fine-tuning, BLASTn [30] is employed with an e-value cutoff of 1e−2 to identify and remove sequences in the pretraining set that are highly similar to fine-tuning sequences. A total of 1,252 sequences is excluded. Although this filtering criterion is conservative, it is adopted to preserve the robustness and validity of downstream model evaluation. Full details of the RNAcentral query procedures and BLAST command-line parameters are provided in **Section S2** of the Supplementary Information.

### 2.2. Pre-training methods

RNA-BERTA is composed of 12 encoder blocks, context length of 512 tokens and an embedding size of 512 (reduced from the standard 768), totaling 55.56M parameters. This architecture is selected to align with the compute–optimal parameter-to-token ratio of 1:20, as proposed by Hoffmann et al. [31]. Tokenization is applied using a character-level Byte-Pair Encoding (BPE) algorithm, rather than a byte-level implementation, to provide a more natural representation of nucleotide tokens. Following the guidelines of Tao et al. [32] and using formulas from approach 1, the vocabulary size is set to 9,700 tokens. Based on this approach, the number of vocabulary parameters is obtained from the compute, which was estimated to be ∼4.05e+17 FLOPS by inverting the formula for the number of non-vocabulary parameters of the model. This estimate closely matches the FLOPs predicted by the Chinchilla framework [31].

Pretraining proceeds in three stages (**Figure 1)**. Because exhaustive hyperparameter optimization (HO) is computationally prohibitive for large transformer models, the μP framework [24] is adapted to enable scalable hyperparameter transferability. In this scheme, the learning rate and number of warm-up steps are optimized on a reduced RNA-BERTa model and then generalized to the full architecture. This strategy improves efficiency by approximately 4.5× compared to direct HO on the complete dataset. HO experiments are executed on four NVIDIA V100 GPUs (150 GB RAM) and completed in ∼48 hours. The complete implementation details are provided in **Section 2.6**. Across all pretraining phases, cosine learning-rate scheduling with warm-up and a 10× decay factor over one epoch is employed, consistent with best practices outlined by Rae et al. [33] and Hoffmann et al. [31]. The μP-compatible implementations of AdamW and RoBERTa [24,34] are used, and attention multiplier is set to √32 to ensure consistent parameter transfer during downstream fine-tuning.

**Figure 1.**
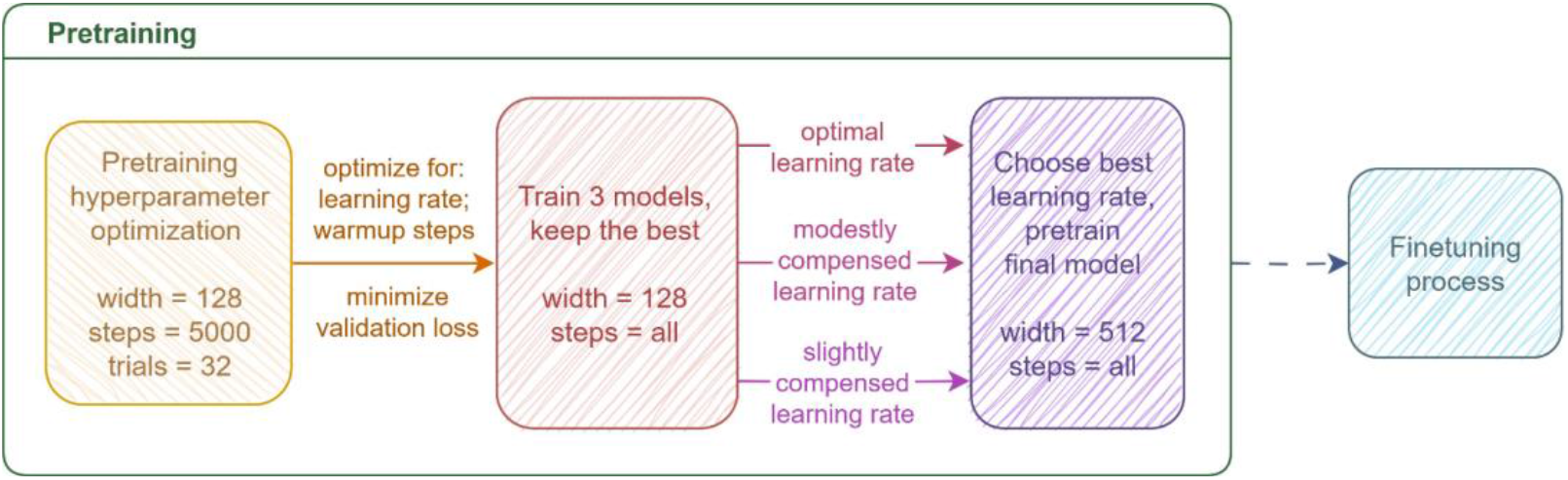
Flow chart of the pretraining hyper parameter selection process, from hyper-parameter optimization to the training of final DLRNA-BERTa model.

### 2.3. Fine-tuning datasets

The fine-tuning dataset comprises 1,439 RNA–drug interaction pairs, each consisting of an RNA sequence, a compound SMILES representation, the dissociation constant (pK_d_), and the corresponding RNA class. The pK_d_ values represent negative logarithm of the dissociation constant (K_d_) values in nano molar concentrations between compound and the RNA target. In total, the dataset includes 759 unique compounds and 294 unique RNA targets. Of 1,439 interaction pairs, 37 RNA sequences exceed the model’s maximum context length (512 tokens) and are truncated. RNA sequence lengths range from 4 to 615 tokens (average: 26.48 tokens), while SMILES strings range from 17 to 333 tokens (average: 70.24 tokens).

This dataset is originally reported by Krishnan et al. [21] and is publicly available in PDF format on the [RSAPred website](https://web.iitm.ac.in/bioinfo2/RSAPred/Predict.html). A minor discrepancy exists between the number of interactions reported in the publication (1,480) and those provided online (1,439). After acquisition, the dataset is parsed and cleaned to remove extraneous columns and ensure compatibility with the model input format. Final datasets are available at: https://huggingface.co/spaces/IlPakoZ/DLRNA-BERTa. Distribution of RNA classes in the fine-tuning dataset is provided in **Table 1**.

**Table 1:** Number of interactions in the training data by RNA class. Values shown in bold indicate class-specific sample counts that differ from those reported in the Krishnan et al [21].

| RNA sequence type | N. of interactions |
| --- | --- |
| Aptamers | **520** |
| miRNAs | 146 |
| Repeats | 97 |
| Ribosomal | **295** |
| Riboswitches | **100** |
| Viral RNAs | **281** |
| **Total** | **1,439** |

### 2.4. Fine tuning methods

The pretrained RNA model is integrated with ChemBERTa [23] to fine-tune seven distinct models. One model is trained on the full set of RNA–drug interactions (the combined model), while the remaining six are trained exclusively for each RNA class. For each model type, 25 hyperparameter optimization trials are conducted using Optuna with the Tree-structured Parzen Estimator (TPE) [35] sampler. The optimized hyperparameters include learning rate, weight decay, dropout, and scheduler decay ratio. Each model is subsequently trained and evaluated using cross-validation (CV) to assess training and validation performance. For the combined model, CV is repeated across three distinct hyperparameter configurations to ensure stability and robustness. To preserve class balance, stratified sampling by RNA class is applied when constructing training and validation splits.

Prior to fine-tuning, dissociation constant (pKd) values are z-score standardized to promote faster and more stable convergence. After CV, the best-performing configuration for each RNA class specific model is selected, while three high-performing configurations are retained for the combined model. All models are then retrained on the full available data for their respective RNA classes, and final performance is reported against the baseline of Krishnan et al. [21]. Detailed evaluation results are presented in **Section 3.3.3**.

### 2.5. External evaluation

All selected models are validated against the external ROBIN classification test datasets provided by the Krishnan et. al. [21]. These datasets contain a total of 5,534 RNA–drug pairs, including 2,538 negative and 2,991 positive interactions. The four datasets used consist of:

- 1,617 aptamer pairs, comprising 5 unique RNA sequences
- 630 miRNA pairs, comprising 3 unique RNA sequences
- 1,459 riboswitch pairs, comprising 5 unique RNA sequences
- 1,828 viral RNA pairs, comprising 6 unique RNA sequences

For the all-types setup, all three models are validated against concatenated test datasets, and the best performing model is selected.

### 2.6. Model architecture for proposed DLRNA-BERTa

A representation of the overall model is shown in **Figure 2**. The model is composed of two transformer encoders:

**Figure 2:**
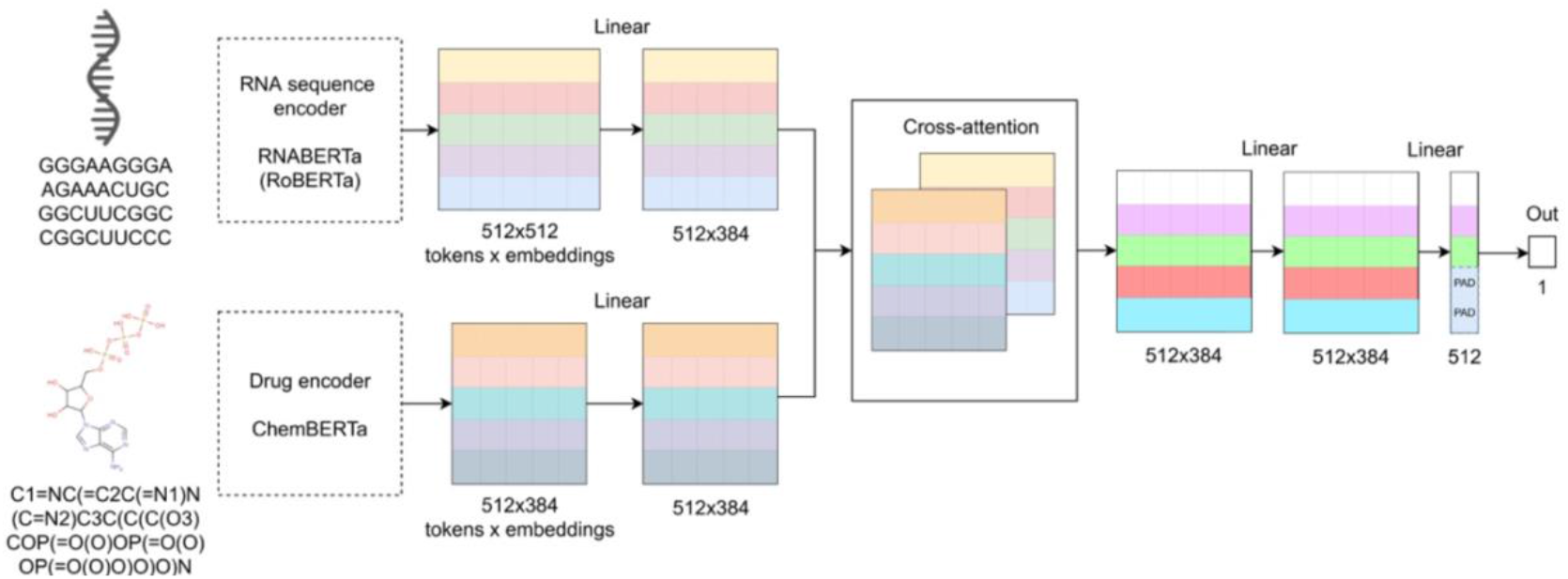
Overall model architecture, encompassing pretrained models for the fine-tuning task.

- RNABERTa, the target encoder, which takes RNA sequences as input and returns sequence embeddings.
- ChemBERTa-v2, the compound encoder, which processes SMILES strings and returns compound embeddings.

ChemBERTa-v2 (77M-MTR) [23] produces embeddings of size 384, whereas RNA-BERTa produces embeddings of size 512. To maintain consistent embedding dimensions, RNA-BERTa outputs are projected to 384 through a linear layer. Both RNA and compound embeddings are then passed through linear layers before being fed into a cross-attention module that serves as the interaction head. This module employs a single attention head to promote interpretable and biologically meaningful embeddings.

Within the cross-attention mechanism, the drug SMILES representation provides the key and value matrices, while the RNA sequence provides the queries matrix. The drug is designated as the key following a lock-and-key analogy, where the RNA sequence is the lock and the drug’s SMILE is the key. The context-updated RNA token embeddings are subsequently passed through a linear integration layer, followed by a second linear layer that reduces the embedding size from 384 to 1. Because linear layers operate independently across embedding dimensions, token identity is preserved. Finally, the token representations are passed through a final linear layer, where vector–vector multiplication is decomposed into two steps: (**i**) element-wise multiplication between input and weight vector components, and (**ii**) summation of the resulting values as shown in following equaton.

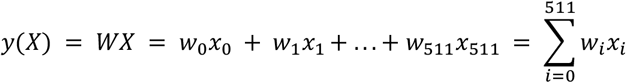

Here, *w*_*i*_*x*_*i*_ represents the contribution of token *i* to the prediction. Prior to summation, a learned padding value replaces contributions for all padding tokens, ensuring equal contribution across them. The summation yields the predicted binding affinity (*pK*_*d*_) between the compound and RNA target. These token-level contributions are further analyzed during interpretation. Both unnormalized and normalized visualization plots are generated, where contributions from padding tokens are redistributed among non-padding tokens. For normalized plots, negative contributions are iteratively reassigned to positive tokens until all values are non-negative. Special tokens such as “<s>“ and “</s>“ are retained during fine-tuning and interpretation to ensure consistency with pretraining process. The cross-attention mechanism also facilitates ablation and perturbation studies, with an example provided in **Section 3.1.2**.

### 2.7. Hyperparameter optimization (HO)

HO for performed for both pretraining and fine-tuning. For pretraining, 32 optuna trials are executed using the TPE sampler. Because *µ*P supports transfer of only non-regularization parameters [24], weight decay is fixed to the standard value of 0.01. Sixteen-bit floating point precision is not used during HO, as it is reported to cause unstable training under µP [36]. For post–layer normalization transformers like RoBERTa [34] implementation, Yang et. al. [24] demonstrate that optimal hyperparameters transfer reliably across different training step counts. Leveraging this, each HO model is trained for 5,000 steps rather than the full 38,000 steps supported by the dataset, thereby reducing computational cost while preserving transferability of the selected hyperparameters. To account for the slight left-shift in the optimal learning rate reported by Yang et al., three additional models are trained with learning rates equal to the HO-derived optimal value multiplied by 1.0, 0.875, and 0.75, respectively.

Fine-tuning HO is conducted using 10-fold CV to account for variability due to training set partitioning. To accelerate optimization, each of the 25 Optuna trials is evaluated on a subset of k = 5 folds out of the 10 total, resulting in 125 models trained per model type, each for 100 epochs. Optimization is performed with the objective function Weighted Mean Squared Error (WMSE), a variant of mean squared error where updates are weighted by a factor proportional to the abundance of the corresponding RNA target class in the dataset. The WMSE is defined as:

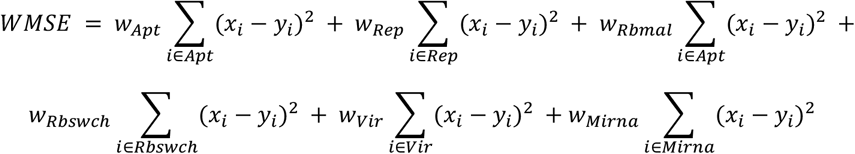

where *w* correspond to class specific weights computed based on class abundance. Exact values are reported in **Section 6** in Supplementary information. WMSE is chosen to promote model generalization for different sequence types, given the different distribution of pK_d_ values by RNA class in the dataset shown by preliminary data analysis (see **Figure 3**). During fine-tuning, we utilized a linear learning rate decay scheduler.

**Figure 3:**
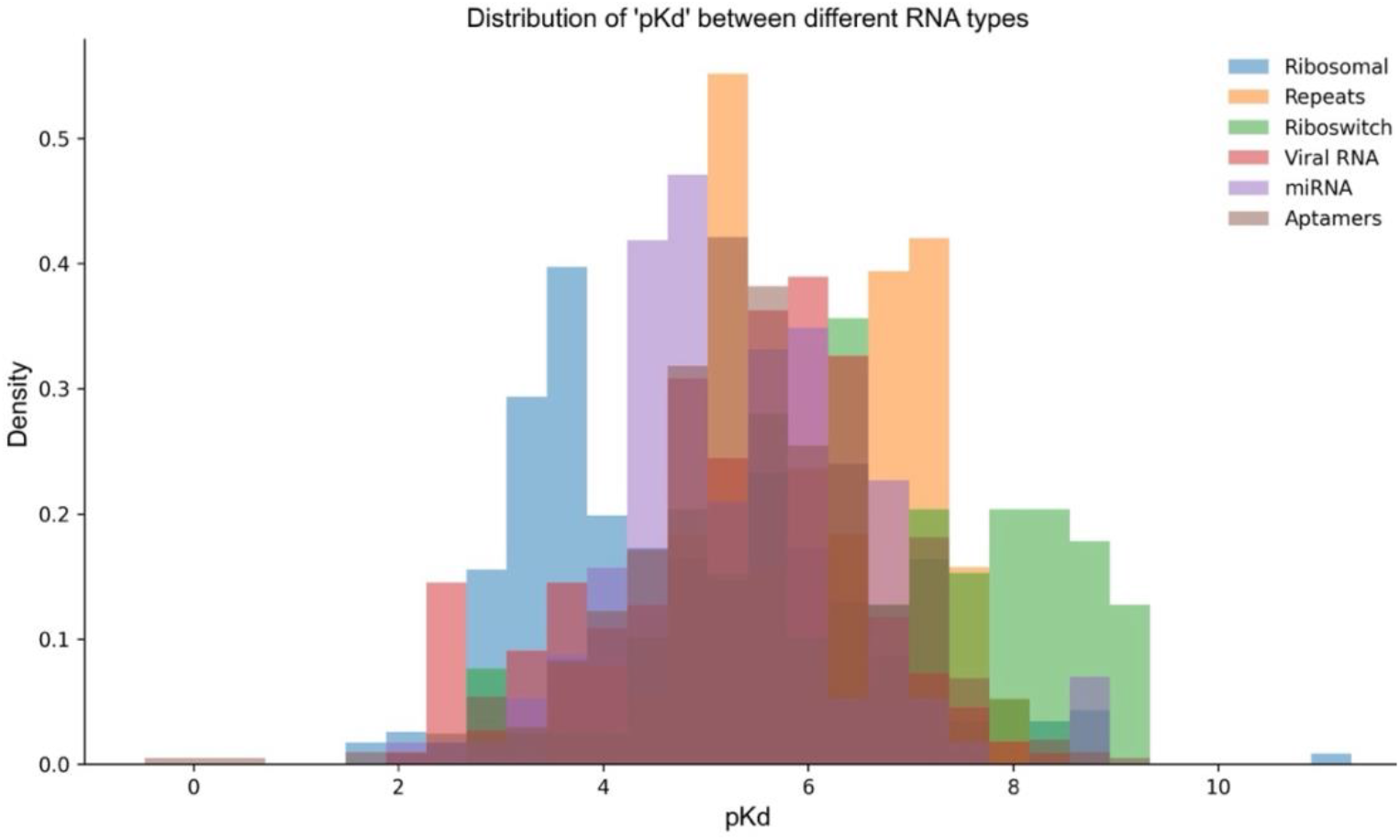
Bar plot of different pKd values by RNA target class.

### 2.8. Evaluation metrics

During pretraining, model quality is evaluated using the bits-per-character (BPC) metric across all phases. BPC is a standard pretraining metric that measures the average number of bits required to encode each character in the text, thereby quantifying how well the model predicts the next character in a sequence.

For fine-tuning HO, the validation set is assessed using three metrics: R^2^ score, mean absolute error (MAE) and pearson’s correlation coefficient (R). The R^2^ metric quantifies the proportion of variance in the dependent variable (pK_d_) explained by the independent variables (RNA sequences and compound SMILES strings). MAE measures the average absolute difference between predicted and actual pK_d_ values. The final model performance is evaluated using R^2^, MAE, and R.

### 2.9. Web application for proposed DLRNA-BERTa

The general model is deployed on a dedicated Hugging Face Space (https://huggingface.co/spaces/IlPakoZ/DLRNA-BERTa) to provide free and user-friendly access. Only the general model is deployed, as the limited training data available for the six class-specific models may restrict their ability to generalize effectively to novel RNA targets submitted by end users. The graphical user interface (GUI) is organized into four tabs: *Prediction & Analysis, Visualizations, Model Settings*, and *About*. By default, users are directed to the *Prediction & Analysis* tab, which provides input fields for an RNA sequence and a compound SMILES string, along with two functional buttons: *Predict Interaction* and *Generate Visualizations*. A snapshot of this feature is shown in **Figure 4**.

**Figure 4:**
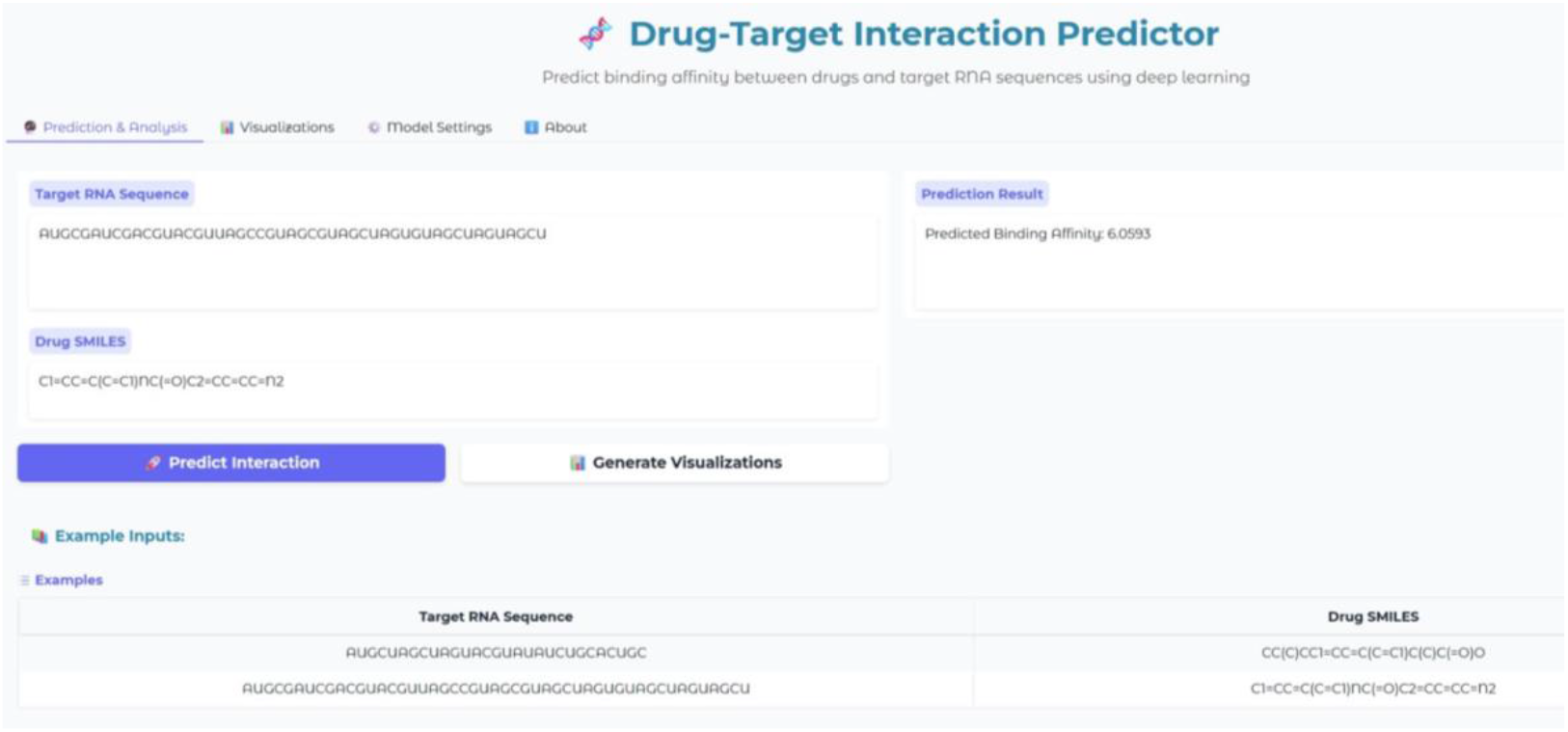
A snapshot of the *Prediction & Analysis* tab on the DLRNA-BERTa web application (https://huggingface.co/spaces/IlPakoZ/DLRNA-BERTa). The predicted binding affinity (pK_d_) is displayed in the *Prediction Result box*.

**Figure 5:**
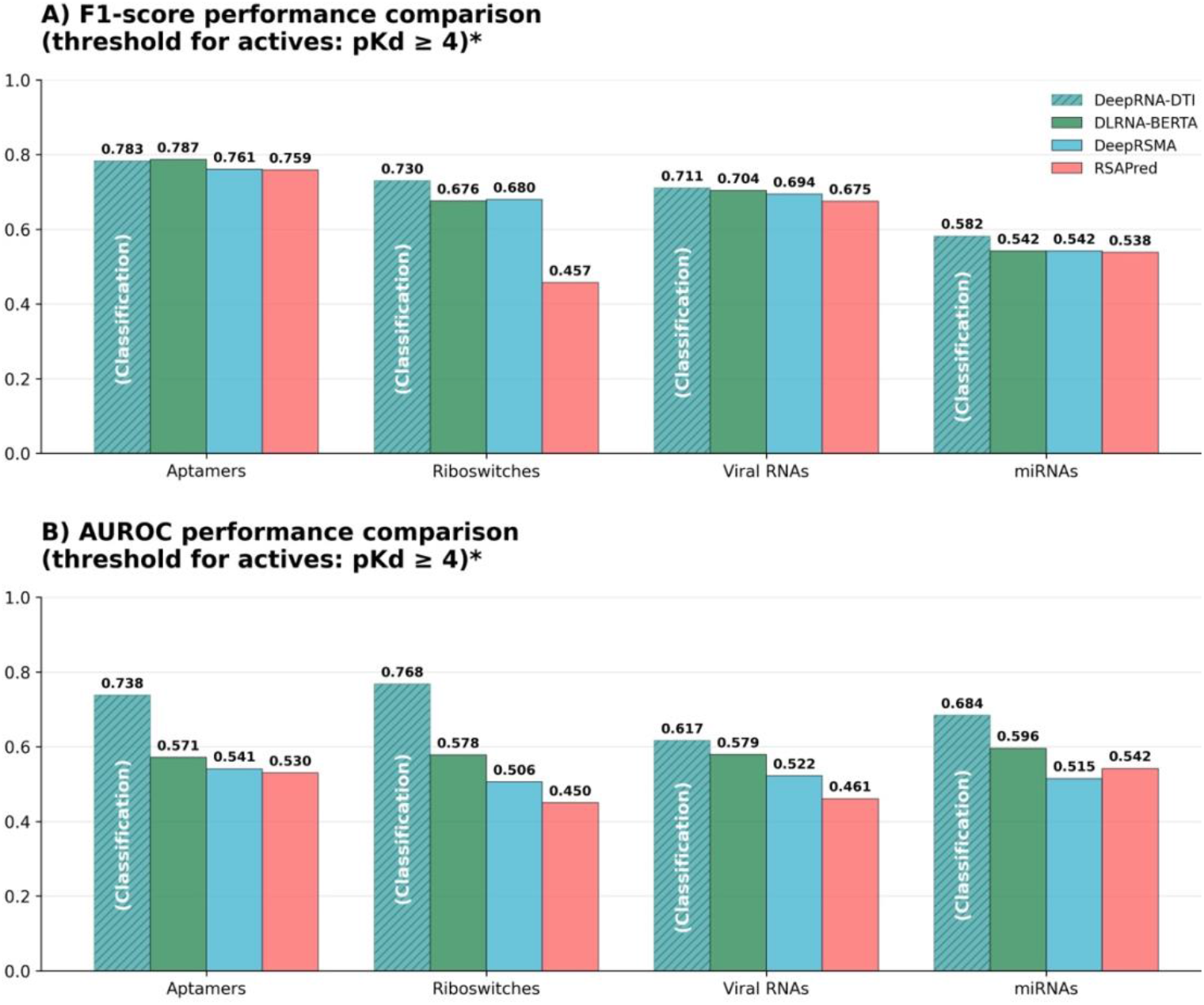
Performance comparison of RNA–drug interaction prediction models across four RNA classes. Bar plots show **(A)** F1-score and **(B)** AUROC performance for DLRNA-BERTA, RSAPred, DeepRNA-DTI, and DeepRSMA models evaluated for aptamers, riboswitches, miRNAs, and viral RNAs classes.DeepRNA-DTI, which employs a classification approach, is indicated by diagonal hatching and labeled “(Classification)” on the corresponding bars. This model achieves higher performance due to greater data availability and a different objective function. Regression-based models were evaluated using a threshold of pKd ≥ 4 to define active compounds. *For the classification model, the threshold that maximized the F1-score was selected.

When the *Predict Interaction* button is clicked, the model computes the predicted pK_d_ value, which is displayed in the *Prediction Result* text box. Alternatively, clicking *Generate Visualizations* provides both the predicted pK_d_ and a set of plots comparable to those presented in this study. These include a cross-attention heatmap, as well as unnormalized and normalized pK_d_ contribution plots. The generated plots are accessible through the Visualizations tab. The Model Settings tab is designed to allow users to select between different model variants; however, at present only the general model is available. The *About* tab provides general information on the model’s usage. At the footer of the GUI, the *Use via API* option provides instructions for programmatic access. Supported languages include Python, JavaScript, and Shell. An example set of Python commands for accessing the API of the proposed DLRNA-BERTa model is provided below.

> from gradio_client import Client
>
> client = Client(“IlPakoZ/DLRNA-BERTa”)
>
> result = client.predict(
>
> target_seq=“AUGCUAGCUAGUACGUAUAUCUGCACUGC”,
>
> drug_smiles=“CC(C)CC1=CC=C(C=C1)C(C)C(=O)O”,
>
> api_name=“/predict_wrapper”
>
> )
>
> print(result)

The *target_seq* field specifies the target RNA sequence, while *drug_smiles* provides the drug’s chemical structure in SMILES format. The *api_name* specifies which API to use, specifically /predict_wrapper is used to make a prediction. The output, stored in the result variable, corresponds to the predicted binding affinity expressed as pK_d_.

## 3. RESULTS AND DISCUSSION

### 3.1. Training and validation performance

#### 3.1.1. Pretraining

Hyperparameter optimization over 5,000 training steps yields an optimal learning rate of 0.00063 and 765 warm-up steps. The optimization landscape is observed to be steep near the optimum, with even small increases in learning rate driving the model toward what appears, based on reduced gradient norms, to be a more stable but suboptimal local solution at approximately 11.4 evaluation BPC. A contour plot of the optimization space illustrating this phenomenon is presented in **Figure S3**, while the slice plot in **Figure S4** further demonstrates the sharp sensitivity of the optimization space to minor learning rate increases.

Following HO, three model variants are trained with learning rates of 0.00063, 0.000552, and 0.000443, each using 765 warm-up steps. Among these, the model with a learning rate of 0.000552 achieves the best performance, with an evaluation BPC of 3.20, compared to 3.32 for the HO-identified optimum. The final model is therefore trained with a learning rate of 0.000552, achieving a final evaluation BPC of 1.78. The corresponding training and evaluation loss curves are shown in Figure S5.

#### 3.1.2. Fine-tuning

HO is first performed for the model trained on all RNA classes. A subset of the best fine-tuning HO trials is shown in **Table 2**. Several technical issues are detected during the initial stages of HO, requiring adjustments to correctly configure the trials; these are discussed in detail in **Supplementary Section S5**. Based on validation R^2^ and MAE, trial 13 emerges as the best-performing run. To ensure robust selection of the final model, three promising trials from cross-validation (CV) are chosen for further evaluation:

**Table 2:** Subset of the best fine-tuning hyperparameter optimization trials. All values correspond to validation metrics. *LR* denotes learning rate, *WD* denotes weight decay, and *MAE diff* denotes the mean difference between training and validation MAE.

| N. | LR | WD | Dropout | | Decay ratio | $R^2$ | MAE | MAE diff |
| --- | --- | --- | --- | --- | --- | --- | --- | --- |
|  |  |  | Hidden | Attention |  |  |  |  |
| 13 | 1.06e-4 | 1.31e-3 | 0.078 | 0.391 | ×3 | 0.870 | 0.307 | 0.076 |
| 20 | 1.02e-4 | 1.67e-3 | 0.082 | 0.425 | ×2 | 0.841 | 0.371 | 0.090 |
| 23 | 8.67e-5 | 2.24e-3 | 0.185 | 0.249 | ×2 | 0.806 | 0.450 | 0.083 |
| 15 | 1.05e-4 | 1.14e-3 | 0.156 | 0.395 | ×3 | 0.804 | 0.369 | 0.079 |
| 19 | 9.96e-5 | 1.71e-3 | 0.166 | 0.345 | ×2 | 0.816 | 0.365 | 0.067 |

- trial 13, corresponding to the best overall validation performance,
- trial 20, selected for competitive metrics, and
- trial 23, included to explore a configuration with a lower learning rate and potentially reduced overfitting.

The hyperparameters and CV results for these three trials are reported in **Table 3**. During this selection, priority is given to higher R^2^ or lower MAE, smaller differences between training and validation MAE, and lower learning rates. For the RNA class–specific models, one set of hyperparameters is selected per class, prioritizing MAE and MAE difference; unlike the combined dataset model, only a single configuration is retained. The results of training for each RNA class are provided in **Table 4**.

**Table 3:**
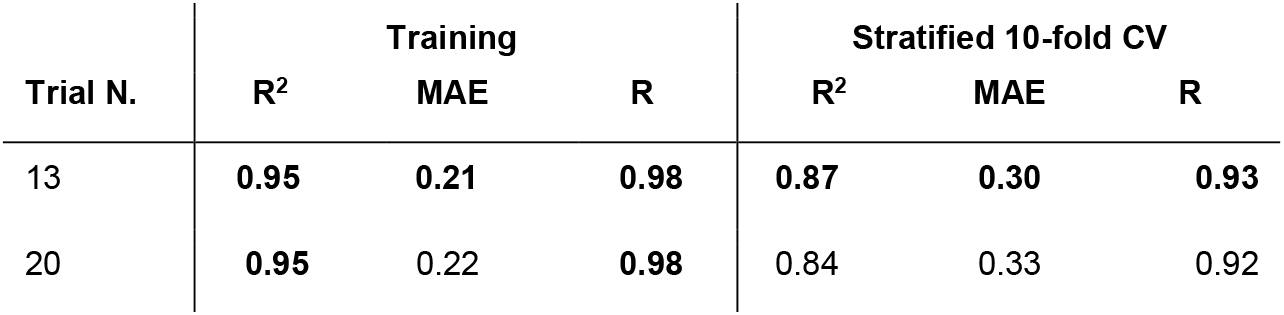
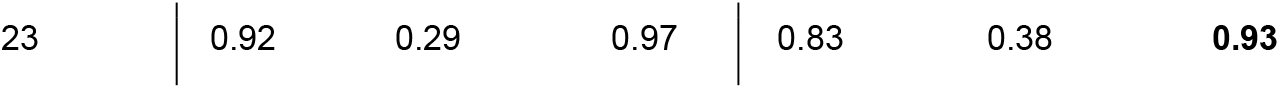
Results of general model training on the three selected trials from HO. In bold, the highest values per column.

| Trial N. | Training |  |  | Stratified 10-fold CV |  |  |
| --- | --- | --- | --- | --- | --- | --- |
| | $R^2$ | MAE | R | $R^2$ | MAE | R |
| 13 | **0.95** | **0.21** | **0.98** | **0.87** | **0.30** | **0.93** |
| 20 | **0.95** | 0.22 | **0.98** | 0.84 | 0.33 | 0.92 |
| 23 | 0.92 | 0.29 | 0.97 | 0.83 | 0.38 | **0.93** |

**Table 4:**
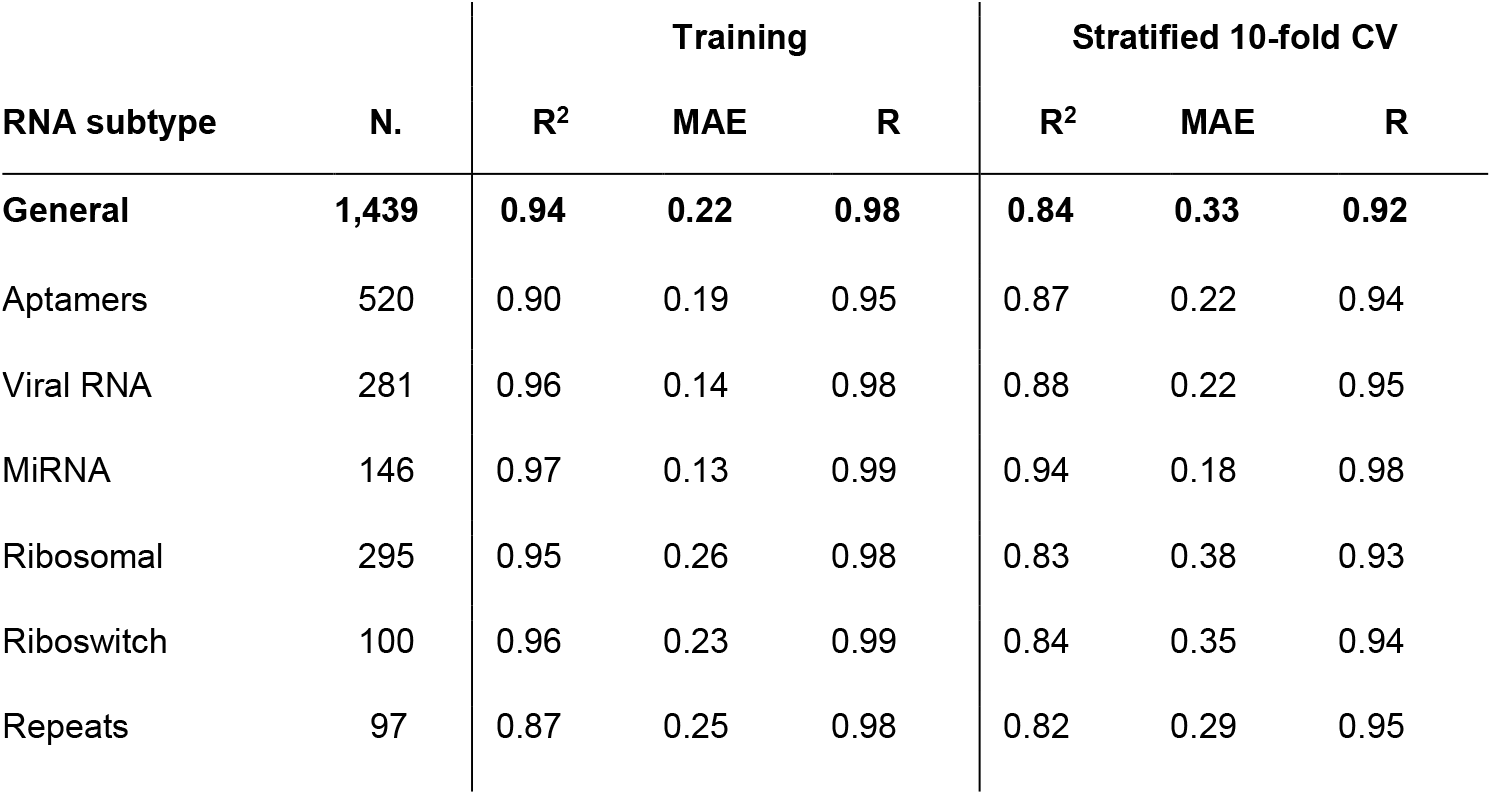
Results of model training on all RNA classes. Column “R” indicates pearson correlation.

Examples of fold-level validation predictions are shown in **Figure S6**. As 10-fold CV is employed, some models have relatively few validation samples per fold, making fold-level performance plots less reliable individually. However, averaging across the 10 folds provides a more stable estimate of true model performance. Notably, Krishnan et al. [21] also adopt 10-fold CV for their baseline models. The final models are retrained on the combined training and validation sets, after which interpretation analyses are performed. From the final linear layer, values and weights are extracted, and both normalized and unnormalized visualizations are generated for all samples with predicted binding affinity pKd > 8 and MAE < 0.5. As an illustrative case, the interaction between the *Bacillus subtilis* flavin mononucleotide (FMN) riboswitch aptamer and its ligand FMN is analyzed (**Figure S7**). In the unnormalized visualization, tokens contribute both positively and negatively to affinity prediction. The strongest positive contributions arise from “<s>“ and “UUCGGGGC,” whereas “UAGUAA” contributes most negatively. This highlights subsequences that most strongly influence predicted function for the given compound, though not necessarily through direct binding. The normalized visualization (**Figure S8**) yields similar results, as negative contributions are minimal and redistributed.

A second example involves **the** bacterial 50S ribosomal RNA subunit and the antibiotic azithromycin (**Figures S9–S10**). Here, differences between normalized and unnormalized visualizations are more pronounced, revealing alternating blocks of positive and negative contributions reminiscent of conservation profiles. We hypothesize that such patterns may reflect functional roles of specific sequence regions in RNA folding and activity. This hypothesis is supported by the consistency of token-importance profiles across different compounds for the same sequence (**Figure S11**), although the limited compound diversity per RNA sequence in the training data may also contribute.

Ablation studies are further performed to probe interpretability. In one case, the FMN riboswitch aptamer with FMN is examined. **Figure S12** shows pre-ablation cross-attention weights, where the ChemBERTa “[CLS]” token exhibits the strongest attention, particularly toward the subsequence “AUUCAG” (weight ≈ 32). When this weight is set to zero, the corresponding cell in the heatmap is removed (**Figure S13**), leading to a slight increase in attention weight for “AUUCAG” but a decrease in predicted pKd, thereby increasing error. In this case, the attention mechanism downweights the subsequence, and its removal redistributes attention via softmax, ultimately reducing affinity prediction accuracy.

Another ablation example examines the interaction between 2-amino-N6-hydroxyadenine and the *Bacillus subtilis* guanine riboswitch (**Figures S14–S15**). Here, the highest weight (≈ 0.18) corresponds to the interaction between subsequence “UGUCAG” and a defined carbon atom of the compound. Setting this weight to zero increases the final token contribution and predicted pKd, again worsening error. As in the FMN case, cross-attention appears to downweight specific subsequences, with removal leading to compensatory redistribution and a shift in predictions.

### 3.2. Comparison with other methods

Training and validation performance is first compared with the baseline models of Krishnan et al. (RSAPred) [21]. Across all RNA classes, the proposed models consistently outperform their results. This improvement reflects the capacity of sufficiently large architectures to capture complex relationships between compound–RNA target pairs, surpassing RSAPred, which relies on a simpler linear regression framework. In addition, the present cross-validation approach does not include stratification by pK_d_, making it more generalizable. Although random splitting with RNA class stratification is not ideal, since RNA targets or compounds may still appear in both training and validation sets, the results nevertheless demonstrate that the proposed models achieve higher performance than RSAPred, even when evaluated on targets or compounds represented in some form within the training set. The obtained results are summarized in **Table 5**.

**Table 5:**
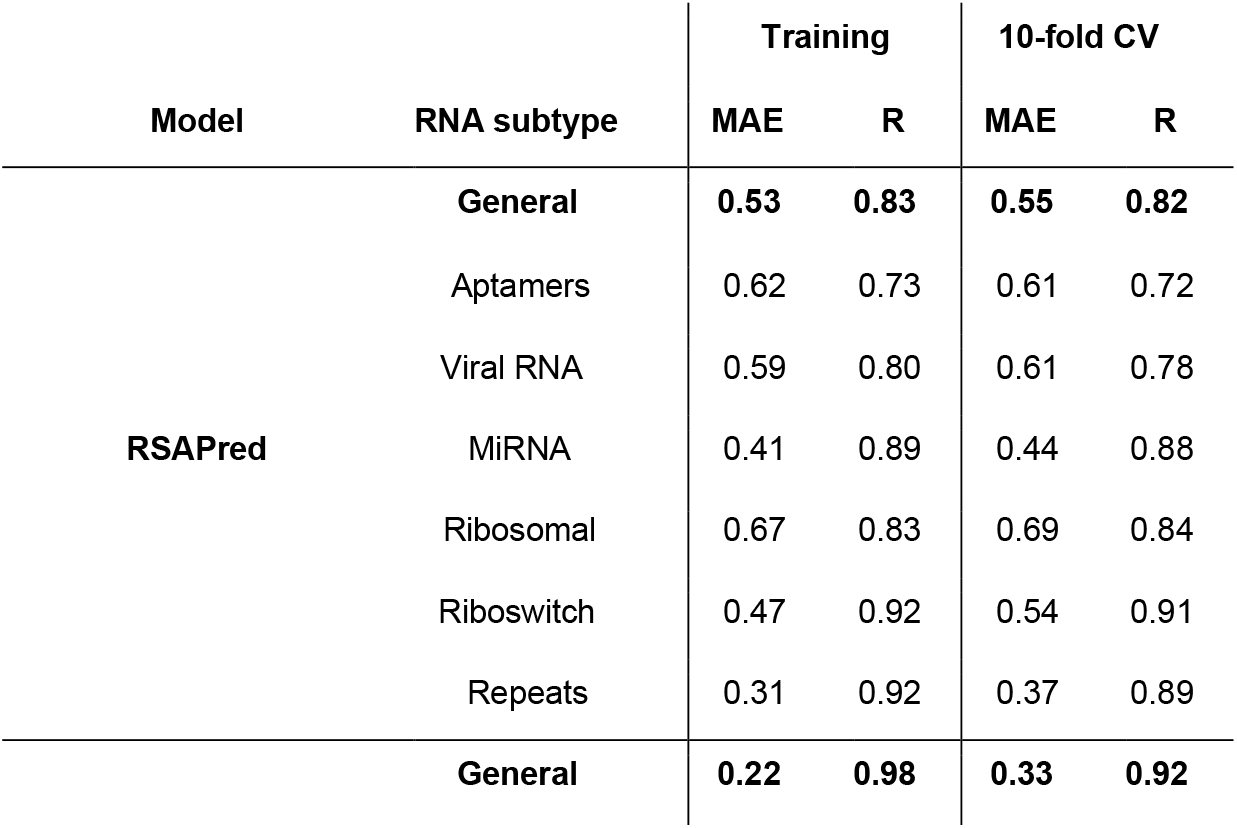
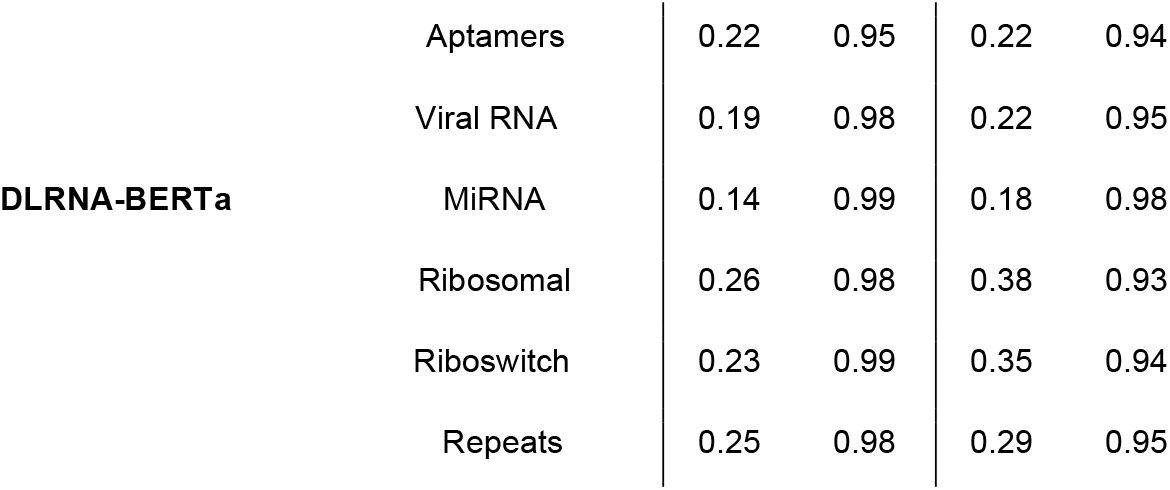
Comparison between training and validation performance of DLRNA-BERTa and RSAPred by Krishnan et al. [21]. R indicates Pearson R correlation.

Performance is next evaluated on the ROBIN datasets provided by Krishnan et al. In this setting, the results obtained are substantially lower than those reported by the authors. For example, Krishnan et al. report an AUROC of 0.971, precision of 0.957, specificity of 0.963, and an F1-score of 0.972 for the aptamer test set, with similarly high performance for miRNAs, riboswitches, and viral RNAs. These values represent remarkably strong predictive performance. Initially, the comparatively lower scores of the present models are attributed to potential overfitting: being simpler, the authors’ regression-based model is less prone to this issue. As a cautionary step, two independent approaches are used to attempt reproduction of the reported results:

- Using the parameters reported in the supplementary table of Krishnan et al., a Python script is implemented to recompute predictions according to the method of Krishnan et al.
- A random sample of 20 positive and 20 negative pairs is drawn from each test dataset, and the authors’ publicly available web interface is queried, followed by a simple statistical analysis of the outputs.

For the first method, all data features are recomputed from scratch and binding affinity is regressed using the parameters as reported in Krishnan et al. [12]. Following the authors’ procedure, classification labels are assigned using a threshold of pKd ≥ 4. Compound features are computed with Mordred [37], a molecular descriptor calculator, and RNA features are extracted using a customized Python script. To validate feature extraction, the computed values are confirmed to match those provided for the nine drug–target pairs available in the GitHub repository of Krishnan et al. Despite this consistency in feature extraction, a substantial discrepancy is observed between the reported performance metrics and those obtained from reproduced predictions. **Table 6** compares the reported and reproduced predictions, while **Table 7** presents the performance of the proposed model alongside the actual performance of the authors’ model under reproduction.

**Table 6:** Expected performance reported by Krishnan et al. [21] compared with performance measured through re-implementation of their model. *Sensit*. and *Specif*. denote sensitivity (recall) and specificity, respectively; *F1* denotes the F1 score.

| RNA classes | Performance reported in RSAPred |  |  |  | Our reproduced performance |  |  |  |
| --- | --- | --- | --- | --- | --- | --- | --- | --- |
|  | Sensit. | Specif. | F1 | AUROC | Sensit. | Specif. | F1 | AUROC |
| Aptamers | 0.98 | 0.96 | 0.97 | 0.97 | 0.98 | 0.02 | 0.76 | 0.53 |
| Riboswitches | 0.97 | 0.94 | 0.96 | 0.95 | 0.44 | 0.47 | 0.46 | 0.45 |
| miRNAs | 0.97 | 0.97 | 0.96 | 0.97 | 0.97 | 0.06 | 0.54 | 0.54 |
| Viral RNA | 0.98 | 0.99 | 0.99 | 0.98 | 0.98 | 0.01 | 0.68 | 0.46 |

**Table 7:** Comparison of RSAPred reproduced performance with proposed DLRNA-BERTa. *‘Sensit*.’ and *‘Specif*.’ refer to sensitivity (or recall) and specificity respectively, and *‘F1’* refers to F1 score.

| RNA classes | RSAPred reproduced performance |  |  |  | DLRNA-BERTa performance |  |  |  |
| --- | --- | --- | --- | --- | --- | --- | --- | --- |
|  | Sensit. | Specif. | F1 | AUROC | Sensit. | Specif. | F1 | AUROC |
| Aptamers | 0.98 | 0.02 | 0.76 | 0.53 | 1.0 | 0.002 | 0.79 | 0.57 |
| Riboswitches | 0.44 | 0.47 | 0.46 | 0.45 | 0.94 | 0.04 | 0.68 | 0.58 |
| miRNAs | 0.97 | 0.05 | 0.54 | 0.54 | 0.99 | 0.04 | 0.54 | 0.60 |
| Viral RNA | 0.98 | 0.01 | 0.68 | 0.46 | 0.94 | 0.1 | 0.70 | 0.58 |

To avoid complications arising from potential inaccuracies in the reported parameters, implementation inconsistencies, or coding errors, the publicly available models hosted on the authors’ website are also evaluated. Because manual entry of every sample is impractical, a random sample of 20 positive and 20 negative labeled examples is drawn with replacement from each dataset and tested against the corresponding models. A simple statistical analysis is then performed. As an illustrative case, the aptamer model is used to demonstrate the statistical procedure; calculations for the remaining models are provided in **Section S1**. Evaluation focuses on specificity and sensitivity, applying a threshold of pKd ≥ 4.0 to distinguish active from inactive interactions, consistent with the authors’ reported approach. The aptamer model is reported to achieve a specificity of 0.96 and a sensitivity of 0.98. A one-sided test is conducted with null and alternative hypotheses defined as follows:

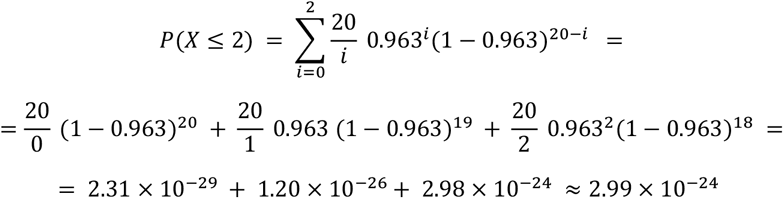

A p-value of 2.99 * 10^−24^ is obtained, indicating an infinitesimal probability that the true specificity is 0.96. Therefore, *H*_0_ is rejected. A similar procedure is applied to sensitivity. We set *H*_0_: *sn* = 0.977 and *H*_*a*_: *sn* < 0.977, and set the significance threshold as before. Again, we computed *P*(*X* ≤ *k*), where this time *k* is the number of correctly predicted positive samples as shown in the following equation. In the aptamers model case, all predicted samples are positive.

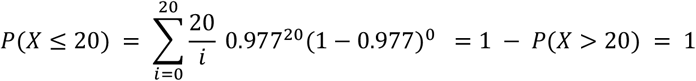

The resulting p-value is 1, we therefore failed to reject the null hypothesis. This indicates that the observed sensitivity is not lower than expected under the reported metric. Together, the results suggest that the model predicts almost exclusively positive labels, yielding very high sensitivity but very low specificity. Consequently, the reported F1 score is likely overly optimistic. These findings align with both the present re-implementation of the authors’ model and the general behavior observed in our own models.

Finally, using the same test data, the proposed model is compared with DeepRSMA by Huang et al. [18], a regression-based approach, and DeepRNA-DTI by Bae et al. [22], a classification-based approach. The proposed model demonstrates superior performance over DeepRSMA and performs competitively with DeepRNA-DTI. Notably, classification models are generally expected to surpass regression models, owing to the broader availability of labeled classification data. The comparative results are presented in **Figure 6**.

**Figure 6:**
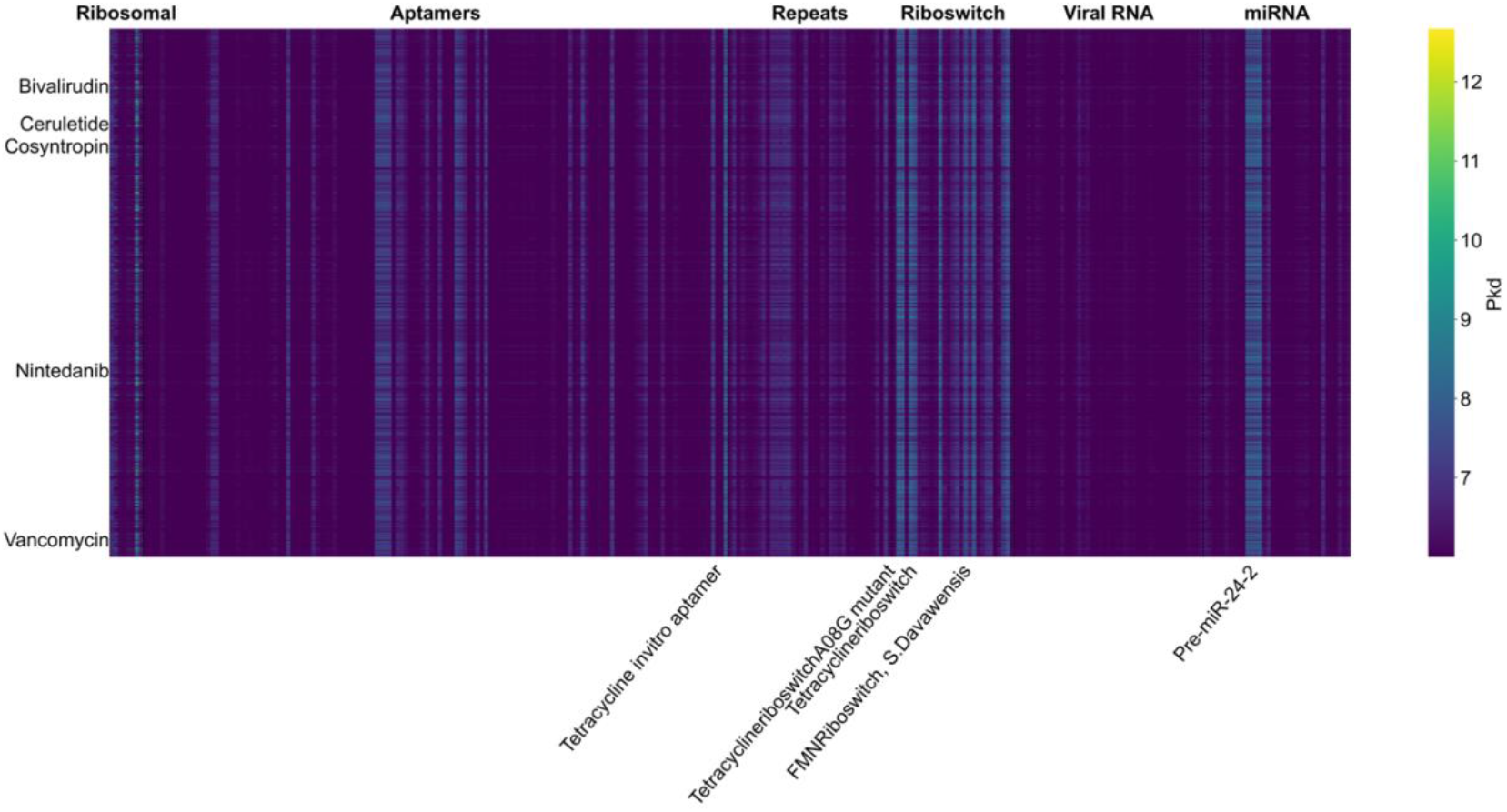
Virtual screening of approved drugs using DLRNA-BERTa. Predicted high-affinity interactions (pK_d_ > 8) are shown across 55 RNA targets, ordered by RNA classes (labelled at top X-axis). The top five drugs with the greatest number of associated targets and the top five RNA targets with the greatest number of associated drugs are highlighted.

### 3.3. Virtual screening of approved drugs against RNA targets

Following a comprehensive evaluation of robustness and predictive performance across both testing and validation datasets, the final models are deployed to predict interactions for each of the six RNA target classes. These models are applied to a dataset of 3,492 approved small molecules sourced from the ChEMBL database (v33). Remarkably, 2,859 compounds are predicted to exhibit strong binding affinities (pKd ≥ 6) across 294 distinct RNA targets, with detailed interaction data provided in **Supplementary File 1**. When the affinity threshold is increased to pK_d_ > 8, 1,613 drugs retain high predicted binding across 55 RNA targets (**Supplementary File 2**). As shown in **Figure 6**, the five drugs with the broadest predicted activity are bivalirudin, nintedanib, vancomycin, ceruletide, and cosyntropin, while the most frequently engaged RNA targets are pre-miR-24-2, petracycline riboswitch A08G mutant, petracycline in vitro aptamer, tetracycline riboswitch, FMN riboswitch, and S. davawensis.

These large-scale predictions highlight opportunities for RNA-targeted drug repurposing by uncovering bioactive off-target interactions among already approved molecules. The resulting hit lists provide a valuable resource for experimental follow-up, guiding biochemical studies to validate predicted RNA binding. As a proof of concept, literature evidence is compiled for bleomycin, one of the 1,695 drugs with predicted high-affinity interactions (pK_d_ > 8), as discussed in **Section 3.4**.

### 3.4. Literature-based validation of bleomycin binding to RNA targets

Our analysis reveals that small molecules predominantly bind to aptamers, viral RNA sequences, repeat elements, and miRNAs, consistent with recent findings reported by multiple groups [38– 43]. miRNAs regulate post-transcriptional gene expression and are implicated in cancer, cardiovascular, metabolic, and neurodegenerative diseases [44,45]. Small molecules capable of binding miRNAs therefore hold substantial therapeutic potential, making this study both translatable and impactful [46]. miRNAs are selected as model RNA targets for small-molecule binding validation in computational pipelines, as they provide several advantages: biological relevance, tractable secondary structures, robust datasets, and experimental feasibility [47]. Their inclusion enhances the credibility, translational potential, and impact of our findings.

Bleomycin, a glycopeptide-derived anticancer agent, emerged as a recurrently predicted RNA-binding candidate by our model, highlighting its consistent interaction with various miRNA and primary transcripts (n=122) (**Supplementary File 1**). Biochemical studies confirm that bleomycin preferentially cleaves purine-rich RNA regions, particularly contiguous AU pairs and RNA hairpins [48], in addition to its well-characterized DNA-binding and cleavage activity [49,50]. Moreover, bleomycin is explored in conjugation with RNA-targeting ligands, enabling dual substrate recognition [51,52]. This design allows the distinct substrate preferences of the RNA-binding ligand and bleomycin to be leveraged for selective degradation of different RNA targets.

Among our predictions, pri-miR-96 (Pri-miR-96RNAC, Pri-miR-96RNA4, Pri-miR-96RNA1, Pri-miR-96RNA5 and Pri-miR-96RNA2), the validated oncogenic miRNAs, emerged as a high-confidence target in our analysis (predicted pK_d_: 6.01, 6.17, 6.23, 6.25 and 6.47 respectively (**Supplementary File 1**). These finding (**Table 8**) align with prior studies showing that Targaprimir-96, a bleomycin conjugate, selectively cleaves pri-miR-96 via sequence-specific recognition [53]. Since multiple individual targets can originate from the same gene family, for example, the pri-miR-96 subfamily includes Pri-miR-96RNAC, Pri-miR-96RNA4, Pri-miR-96RNA1, and Pri-miR-96RNA5, we reported a range of predicted pK_d_ values rather than a single value in **Table 8**.

**Table 8.** Predicted binding affinities of bleomycin (and conjugates) to RNA targets, supported by validation from prior literature.

| RNA | RNA Form | Predicted $pK_d$ | Notes | Ref |
| --- | --- | --- | --- | --- |
| pri-miR-96 | Drosha processing site, UU, GG loop | 6.01 - 6.47 | Accessible/cleavage through conjugation of active compound (Targaprimir-96) | [53] |
| miR-21 | WT | 6.79 | Preferentially to the pre-miRNA stem-loop | [50] |
| miR-17/18a/20a | pre-/ Dicer site/ | 7.17 | Members of miR-17~92 cluster; accessible/cleavage through conjugation of active compound | [50,54] |
| (CCUG)10 | Repeat elements | 6.42 | Accessible/cleavage through conjugation of active compounds | [55] |
| (CUG) 2,4,6, 10 and 16 | Repeat elements | 6.0 - 7.1 | Accessible/cleavage through conjugation of active compounds | [56] |

Additional predicted interactions include miR-21 (pK_d_: 6.79) **and** pre-miR-17/18a/20a (pK_d_: 7.17), components of the oncogenic miR-17-92 cluster, suggesting that bleomycin or its derivatives may exert multi-miRNA regulatory effects through RNA backbone scission or conformational stabilization [50,54] (**Table 8**). Structured repeat elements, including CCUG, (predicted pK_d_: 6.42) and CUG, (predicted pK_d_: 6.0-7.1), are also predicted as bleomycin-binding agents. These interactions are directly relevant to therapeutic strategies targeting myotonic dystrophy types 1 and 2 (DM1 and DM2), where CCUG/CUG repeat expansions are pathogenic [55,56] (**Table 8, Supplementary File 1**).

Collectively, these predicted interactions illustrate the diverse regulatory potential of small molecules on RNA function, including mechanisms such as direct cleavage (in conjugated forms) or allosteric interference with miRNA biogenesis. While bleomycin is presented here as an illustrative case, the broader validation of predicted RNA targets against published literature underscores the strength of the AI-driven approach and its promise for advancing RNA-targeted therapeutic discovery.

## 4. CONCLUSION

In this study, DLRNA-BERTa is introduced, a transformer-based deep learning architecture tailored for predicting binding affinities (pKd) between small molecules and diverse RNA targets using only primary representations (RNA sequences and SMILES strings). The model leverages a cross-attention mechanism to capture contextual interactions between RNA and drug tokens, enabling end-to-end, structure-free prediction of RNA–drug interactions.

Through extensive benchmarking, DLRNA-BERTa demonstrated superior performance compared to existing models. On the general RNA interaction dataset, our model achieved a pearson correlation coefficient of 0.92, R^2^ of 0.84, and mean absolute error (MAE) of 0.33 during 10-fold CV, outperforming the reference model of Krishnan et al. (pearson correlation ≈ 0.82). DLRNA-BERTa maintained strong predictive power across individual RNA subclasses as well, achieving Pearson R values ranging from 0.93 to 0.98, with the highest accuracy observed for viral RNAs and miRNAs. Importantly, even for RNA classes with limited training data, such as repeats and riboswitches, DLRNA-BERTa achieved robust results (e.g., R = 0.95 for repeats, R^2^ = 0.84 for riboswitches), highlighting the model’s generalizability and resilience to class imbalance. Evaluation on external datasets further confirmed the superiority of DLRNA-BERTa over existing regression-based approaches. For instance, on the miRNA subset, our model achieved an AUROC of 0.60, outperforming DeepRSNA (0.52) and RSAPred (0.54). Similarly, DLRNA-BERTa obtained an F1-score of 0.79 on aptamers, exceeding the 0.76 reported by both DeepRSNA and RSAPred.

Application of proposed model to 3,492 approved small molecules from the ChEMBL database identifies 3,223 candidates with predicted binding affinities of pK_d_ ≥ 6 against 278 unique RNA targets. These findings open new opportunities for RNA-targeted drug repurposing, particularly for rare diseases and RNA elements traditionally considered “undruggable.” As proof of concept, literature validation of predicted interactions for bleomycin confirms binding to several oncogenic miRNAs and structured repeat elements, underscoring the biological significance and translational potential of the framework.

Looking forward, future work may improve the current architecture by incorporating geometric deep learning methods, which can better capture structural features of RNA–ligand interactions. The current approach can also be extended to additional ligand classes, such as peptides and oligonucleotides, as suitable training datasets become available. Moreover, the in-silico hit lists generated in this study represent a valuable resource for experimental biochemists, providing guidance for laboratory-based target validation and lead optimization. Finally, the DLRNA-BERTa source code is released under FAIR principles, ensuring accessibility for future researchers to extend the framework and advance RNA-targeted drug discovery.

## ACKNOWLEDGMENTS

The work was funded by the Research Council of Finland (No. 351507 to ZT).

