## Supplementary information for "DLRNA-BERTa: A transformer approach for RNA-drug binding affinity prediction"

#### S1. Statistical tests for the other Krishnan et. al. models

In this supplementary section, we report statistical tests (like the ones employed in **Section 3.2**) for the other RSAPred models. We test the accuracy of the reported results for specificity and sensitivity metrics.

#### S1.1 Riboswitches

Specificity = 0.936 - 14 correctly predicted as negative out of 20.

$$P(X \leq 14) \ = \sum_{i=0}^{14} \frac{20}{i} \ 0.936^{i}(1-0.936)^{20-i} \ = \ 1 - P(X > 14) \ =$$

$$= 1 - \frac{20}{15} 0.936^{15} \ (1-0.936)^{5} \ - \frac{20}{16} 0.936^{16} \ (1-0.936)^{4} \ - \frac{20}{17} \ 0.963^{17}(1-0.963)^{3}$$

$$- \frac{20}{18} \ 0.963^{18}(1-0.963)^{2} \ - \frac{20}{19} \ 0.963^{19}(1-0.963)^{1} \ - \frac{20}{20} \ 0.963^{20} \ \approx$$

$$\approx 1 \ - \ 0.006 \ - 0.028 \ - \ 0.097 \ - \ 0.236 \ - \ 0.364 \ - \ 0.266 \ \approx 0.003$$

Null hypothesis is rejected with p-value $0.003$.

Sensitivity = 0.971 - 8 correctly predicted as positive out of 20.

$$P(X \leq 8) \ = \sum_{i=0}^{8} \frac{20}{i} \ 0.971^{i}(1-0.971)^{20-i} \ =$$

$$= \frac{20}{0}(1-0.971)^{20} + \frac{20}{1} 0.971^{1} \ (1-0.971)^{19} + \frac{20}{2} \ 0.971^{2}(1-0.971)^{18} \ +$$

$$+ \frac{20}{3} 0.971^{3} \ (1-0.971)^{17} + \frac{20}{4} 0.971^{4} \ (1-0.971)^{16} + \frac{20}{5} 0.971^{5} \ (1-0.971)^{15} \ +$$

$$\frac{20}{6} 0.971^{6} \ (1-0.971)^{14} + \frac{20}{7} 0.971^{7} \ (1-0.971)^{13} + \frac{20}{8} 0.971^{8} \ (1-0.971)^{12} \ \approx$$

$$\approx 1.77 \times 10^{-31} \ + \ 1.18 \times 10^{-28} \ + \ 3.77 \times 10^{-26} \ + \ 7.57 \times 10^{-24} + 1.08 \ \times 10^{-21} \ +$$

$$+ 1.15 \times 10^{-19} + 9.67 \ \times 10^{-18} \ + \ 6.47 \ \times 10^{-16} \ + \ 3.52 \ \times 10^{-14} \ \approx 3.588 \ \times 10^{-14}$$

Null hypothesis is rejected with p-value $3.588 \times 10^{-14}$.

**S1.2 Viral RNAs**

Specificity = 0.985 - 2 correctly predicted as negative out of 20.

$$P(X \leq 2) \ = \sum_{i=0}^{2} \frac{20}{i} \ 0.985^{i}(1-0.985)^{20-i} \ =$$

$$= \frac{20}{0} \ (1-0.985)^{20} \ + \frac{20}{1} \ 0.985 \ (1-0.985)^{19} \ + \frac{20}{2} \ 0.985^{2}(1-0.985)^{18} \ =$$

$$\approx 3.33 \times 10^{-37} \ + 4.37 \times 10^{-34} \ + 2.72 \times 10^{-31} \ \approx \ 2.72 \times 10^{-31}$$

Null hypothesis is rejected with p-value $2.72 \times 10^{-31}$.

Sensitivity = 0.983 - 17 correctly predicted as positive out of 20.

$$P(X \leq 17) \ = 1 - \ P(X > 17) =$$

$$= 1 - \frac{20}{18} 0.983^{18}(1-0.983)^{2} \ - \frac{20}{19} \ 0.983^{19} \ (1-0.983)^{1} \ - \frac{20}{20} \ 0.983^{20} \ =$$

$$\approx 1 - \ 0.04 \ - \ 0.25 \ - 0.71 \ \approx \ 0.005$$

Null hypothesis is rejected with p-value $0.005$.

**S1.3 miRNAs**

Specificity = 0.967 - 2 correctly predicted as negative out of 20.

$$P(X \leq 2) \ = \sum_{i=0}^{2} \frac{20}{i} \ 0.967^{i}(1-0.967)^{20-i} \ =$$

$$= \frac{20}{0} \ (1-0.967)^{20} \ + \frac{20}{1} \ 0.967 \ (1-0.967)^{19} \ + \frac{20}{2} \ 0.967^{2}(1-0.967)^{18} \ =$$

$$\approx 2.35 \times \ 10^{-30} \ + 1.37 \times 10^{-27} + 3.83 \times 10^{-25} \approx \ 3.84 \times 10^{-25}$$

Null hypothesis is rejected with p-value $3.84 \times 10^{-25}$.

Sensitivity = 0.966 - 17 correctly predicted as positive out of 20.

$$P(X \leq 20) \ = 1 - \ P(X > 17) = \ 1$$

Null hypothesis is not rejected (p-value 1).

### S2. RNACentral and blast queries

**miRNAs**: ((RNA* AND length:[10 TO 2000] AND (rna_type:"miRNA" OR rna_type:"pre miRNA" )) AND entry_type:"Sequence")

**repeats**: ((repeat* AND length:[10 TO 2000] AND entry_type:"Sequence"))

**ribozymes**: (RNA* AND rna_type:"ribozyme" AND entry_type:"Sequence" AND length:[10 TO 2000]))

**riboswitches**: ((RNA* AND so_rna_type_name:"Riboswitch" AND length:[10 TO 2000] AND entry_type:"Sequence"))

#### **blast queries**:

```
makeblastdb -in database.fasta -input_type fasta -dbtype nucl
blastn -db database.fasta -query to_search.txt -out blast_results.out -outfmt 6 -evalue 1e-2
```

### S3. Optimal RNA-BERTa size calculations

The RNA-BERTa model is composed of 50,952,000 non-vocabulary parameters, number obtained experimentally using a python script, given model hidden size of 512 and 12 encoder blocks. As mentioned earlier, we use approach 1 by Tao et. al. [32] and the specified formulae to calculate the optimal vocabulary size. The formula for calculating the number of non-vocabulary parameters is given by:

$$N_{nv} = 0.08 \times C^{0.50}$$

where C is the compute (in FLOPS). By inverting the formula, we can calculate C:

$$C = \frac{{N_{nv}}^2}{0.08} = (\frac{50{,}952{,}000}{0.08})^2 \approx 4.06 \times 10^{17}$$

Now, we can obtain the number of vocabulary parameters using the second formula and the compute just calculated:

$$N_v = 0.20C^{0.42} = 0.20 \times (4.06 \times 10^{17})^{0.42} \approx 4{,}972{,}913$$

The number of vocabulary parameters in a transformer model is usually the number of parameters of the matrix that assigns to each token its own embedding, which has size $V \times d$, where $V$ is the vocabulary size and $d$ is the hidden size.

So, dividing the number of vocabulary parameters by the hidden size $d = 512$ of our model, we obtain:

$$\frac{4{,}972{,}913}{512} \approx 9712$$

meaning that the optimal vocabulary size, according to approach one of this paper, is 9712 tokens. We approximate to 9700 to get a more appealing number.

#### S4. Masking of the output of cross-attention

To avoid padding tokens creating noise in predictions, the tokens corresponding to padding tokens resulting from cross-attention are manually zeroed out. The attention formula is the following:

$$softmax(\frac{QK^T}{\sqrt{d_k}})$$

The attention matrix $A = \frac{QK^T}{\sqrt{d_K}}$ is the result of matrix-matrix multiplication which results

in queries to be distributed vertically and keys horizontally,

$$\begin{bmatrix} q_{11} & q_{12} & q_{13} \\ & \vdots & \\ q_{n1} & q_{n2} & q_{n3} \end{bmatrix} \times \begin{bmatrix} k_{11} & & k_{l1} \\ k_{12} & \dots & k_{l2} \\ k_{13} & & k_{l3} \end{bmatrix} = \begin{bmatrix} q_1 k_1 & \cdots & q_1 k_l \\ \vdots & \ddots & \vdots \\ q_n k_1 & \cdots & q_n k_l \end{bmatrix}$$

where *n* and *l* correspond to the number of query and key tokens respectfully. Each row corresponds to attention values for one query, and each column corresponds to the attention values for one key. A key padding mask is applied to avoid attention scores to be computed using padding tokens in the key, effectively zeroing out some columns of this matrix. The resulting attention matrix is then multiplied by $V^T$ , resulting in:

$$\begin{bmatrix} a_{11} & \cdots & a_{1l} \\ \vdots & \ddots & \vdots \\ a_{n1} & \cdots & a_{nl} \end{bmatrix} \times \begin{bmatrix} v_{11} & v_{12} & v_{13} \\ & \vdots & \\ v_{l1} & v_{l2} & v_{l3} \end{bmatrix} = \begin{bmatrix} a_1 v_1 & \cdots & a_1 v_l \\ \vdots & \ddots & \vdots \\ a_n v_1 & \cdots & a_n v_l \end{bmatrix} = \frac{1}{\sqrt{d_K}} \begin{bmatrix} q_1 k_1 v_1 & \cdots & q_1 k_l v_l \\ \vdots & \ddots & \vdots \\ q_n k_1 v_1 & \cdots & q_n k_l v_l \end{bmatrix}$$

As before, each row corresponds to one query. Since we already masked out padding key tokens, we only have to mask out queries to avoid padding tokens influencing the results. To do this, we can zero out rows corresponding to each padding token in the queries, similarly to how we masked columns for the keys.

#### S5. Variance collapse

When fine-tuning the model, we observed something that we previously referred to as *variance collapse*. This manifested as a sudden drop in performance in mid-training or failure in improvement of performance after the first epochs and almost perfect convergence to the same value (often close to the mean) in the outputs of the last layers.

This problem is easily detectable through both training and validation metrics, as it manifests itself as a sudden drop in the $R^2$ score to a near-zero value, usually preceded by a short performance instability period. While total collapse was mostly fixed through keeping the learning rate low enough, different versions of this problem presented themselves during the training of the model.

One variation of this problem manifested as a collapse in the performance of the model only when it was set to *eval()* mode (HuggingFace models' mode for evaluation). This version of collapse would not totally reset the performance of the model, but greatly degrade it. Comparing plots of predicted against actual pKd in both *train()* and *eval()* mode, vertical patterns of points were immediately visible in *eval()* mode but not in *train()* mode. **Figure S1** showcases this problem.

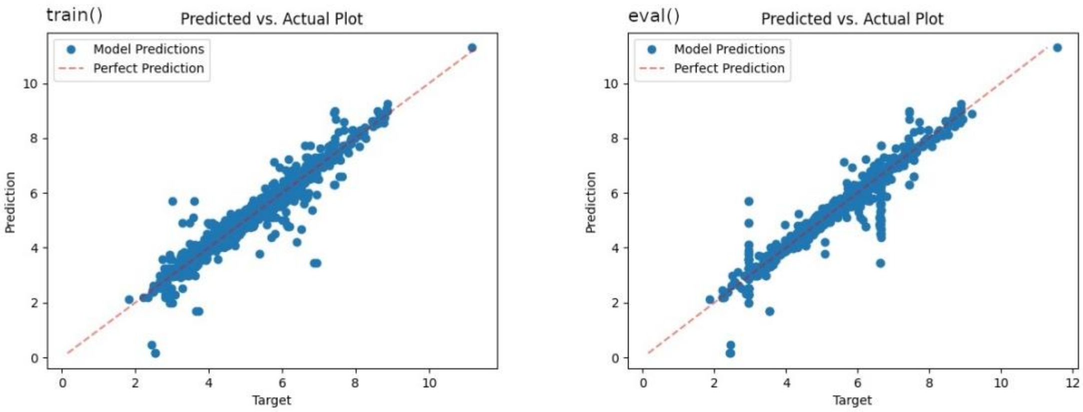


**Figure S1:** This figure shows prediction of the same (training) data using the same model, with the only difference in model mode. A clear vertical pattern of points, otherwise absent, were present when the predictions were made with the model in *eval()* mode.

This problem was not very consistent and manifested only so often. Changing from *train()* to *eval()* mode mainly causes a difference in the behavior of Dropout and BatchNorm layers. Given that our model only utilized the former, we decided to make some tests with dropout disabled: this indeed caused the problem to disappear, causing the performances between the two modes to match. The easiest but maybe not optimal solution to this problem is therefore avoiding the use of dropout.

Looking online, we were able to find this GitHub issue that seemed to resemble our problem. The problem emerges from the use of long type masks, instead of float or bool masks, as padding masks for the TransformerEncoder class, used by RoBERTa models. This did not raise any exception but caused differences in behavior of certain methods, resulting in lower performance. This problem seems to be resolved in newer versions of PyTorch, but not in version 2.5.1, the one we were using.

While after applying this fix the problem seemed to be solved for the general model, variance collapse still happened in the specialized models, but in a slightly different way. The problem is still characterized by a difference between model mode performance and is still caused by dropout. This time, hidden layers' dropout specifically seemed to be the cause of this behavior. A showcase of the problem is shown in **Figure S2**.

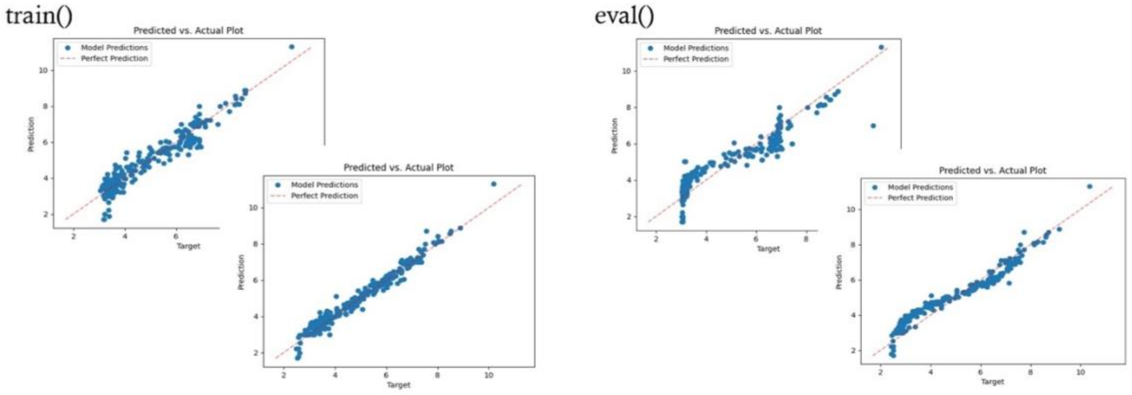


**Figure S2:** This figure shows two examples of predictions with the same (training) data using the same model, with the only difference in model mode. Other than a vertical pattern of points, the *eval()* plot also presents a slightly undulating character in the distribution of points, which is absent in *train()* mode.

The specific causes of this problem are still largely unknown, and attempts to solve it have been largely unsuccessful. However, since this problem can be detected using the training set alone and comparing the performance between *train()* and *eval()* modes, the parameters can be adjusted or the training can be repeated until this problem disappears when the final model is produced. This is not realistic for cross-validation performance evaluation, but that only means that cross-validation performance is likely underestimated.

#### S6. WMSE weights values

Values for the WMSE weights are reported in the table below.

| Parameter | RNA class | Value |
|---|---|---|
| $w_{Apt}$ | Aptamer | 0.46118234 |
| $w_{Rep}$ | Repeat | 2.48084291 |
| $w_{Rbmal}$ | Ribosomal | 0.81140351 |
| $w_{Rbswch}$ | Riboswitches | 2.39814815 |
| $w_{Vir}$ | Viral | 0.85309618 |
| $w_{Mirna}$ | MiRNA | 1.6475827 |

#### S7. Other supplementary figures

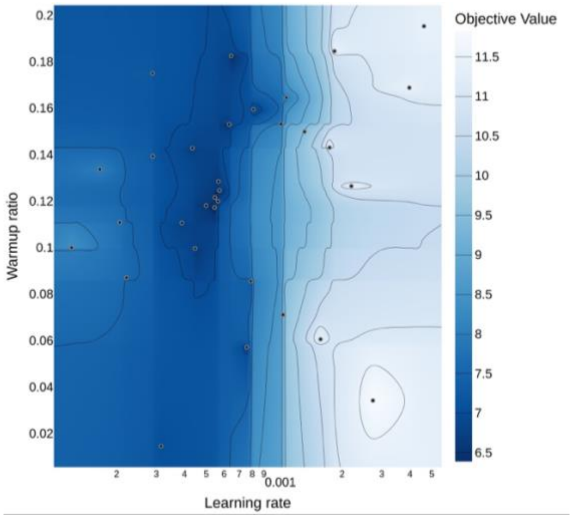


**Figure S3:** Contour plot of the optimization landscape. The x-axis represents variation in learning rate, and the y-axis represents variation in the warm-up ratio (relative to a total of 5,000 steps). A narrow vertical boundary is observed, separating the global optimum region (left, dark blue) from a suboptimal local solution (right, light blue).

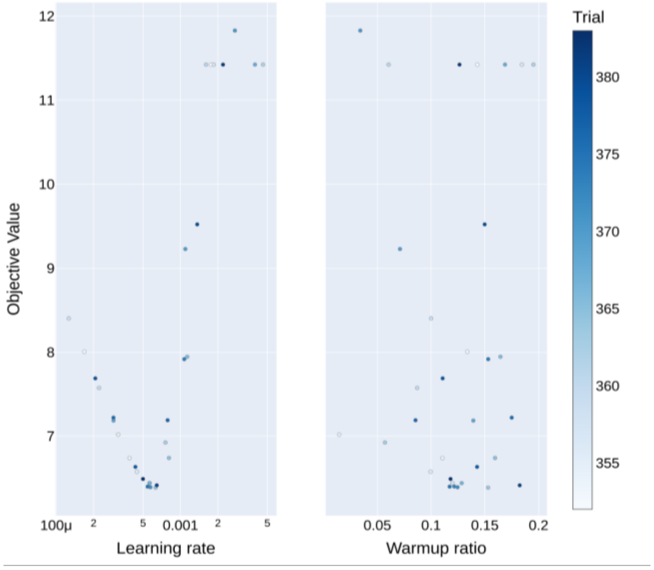


**Figure S4:** Slice plot of learning rate and warm-up ratio (relative to a total of 5,000 steps). In the left panel, the steep slope near the upper bound is evident, as the objective value rises sharply with small increases in learning rate. By contrast, the warm-up ratio shows no clear or consistent correlation with the objective value.

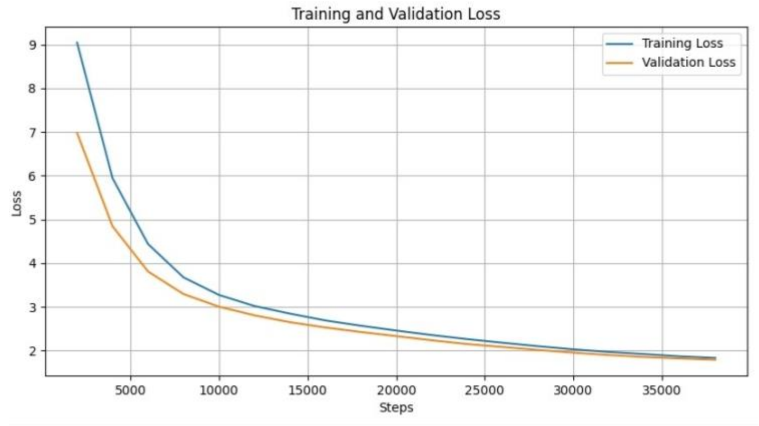


**Figure S5:** Evaluation and training loss curves for the final model pretraining.

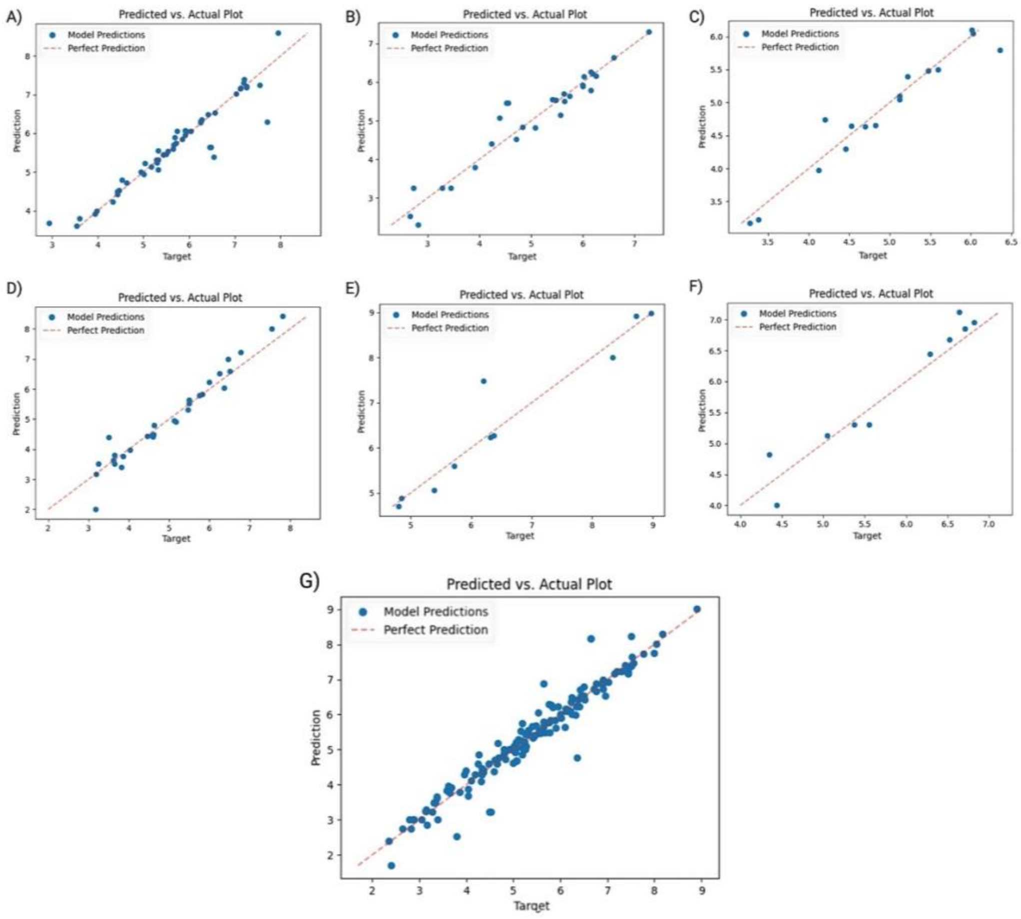


**Figure S6:** Visualization of predicted versus actual pKd values for one validation split from each model. Model types are: **(A)** Aptamers, **(B)** Viral RNA, **(C)** miRNA, **(D)** Ribosomal RNA, **(E)** Riboswitch, **(F)** Repeats, and **(G)** General model encompassing all RNA classes.

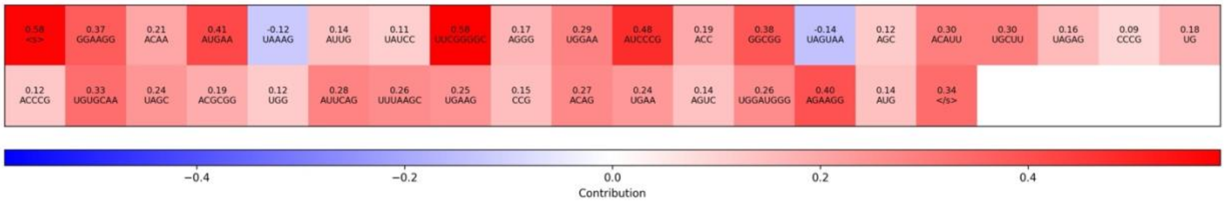

**Figure S7:** Unnormalized visualization between target “Bacillus subtilis’s FMN riboswitch aptamer” and compound “FMN”. In each cell, the predicted binding affinity contribution for each token and the sequence corresponding to that token can be positive, zero or negative.

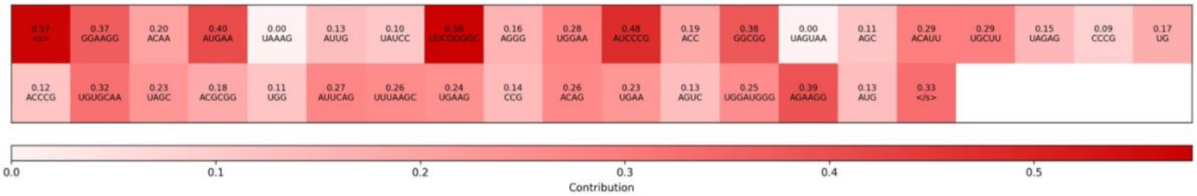

**Figure S8:** Normalized visualization between target “Bacillus subtilis’s FMN riboswitch aptamer” and compound “FMN”. In each cell, the predicted binding affinity contribution for each token and the sequence corresponding to that token can be only positive or zero.

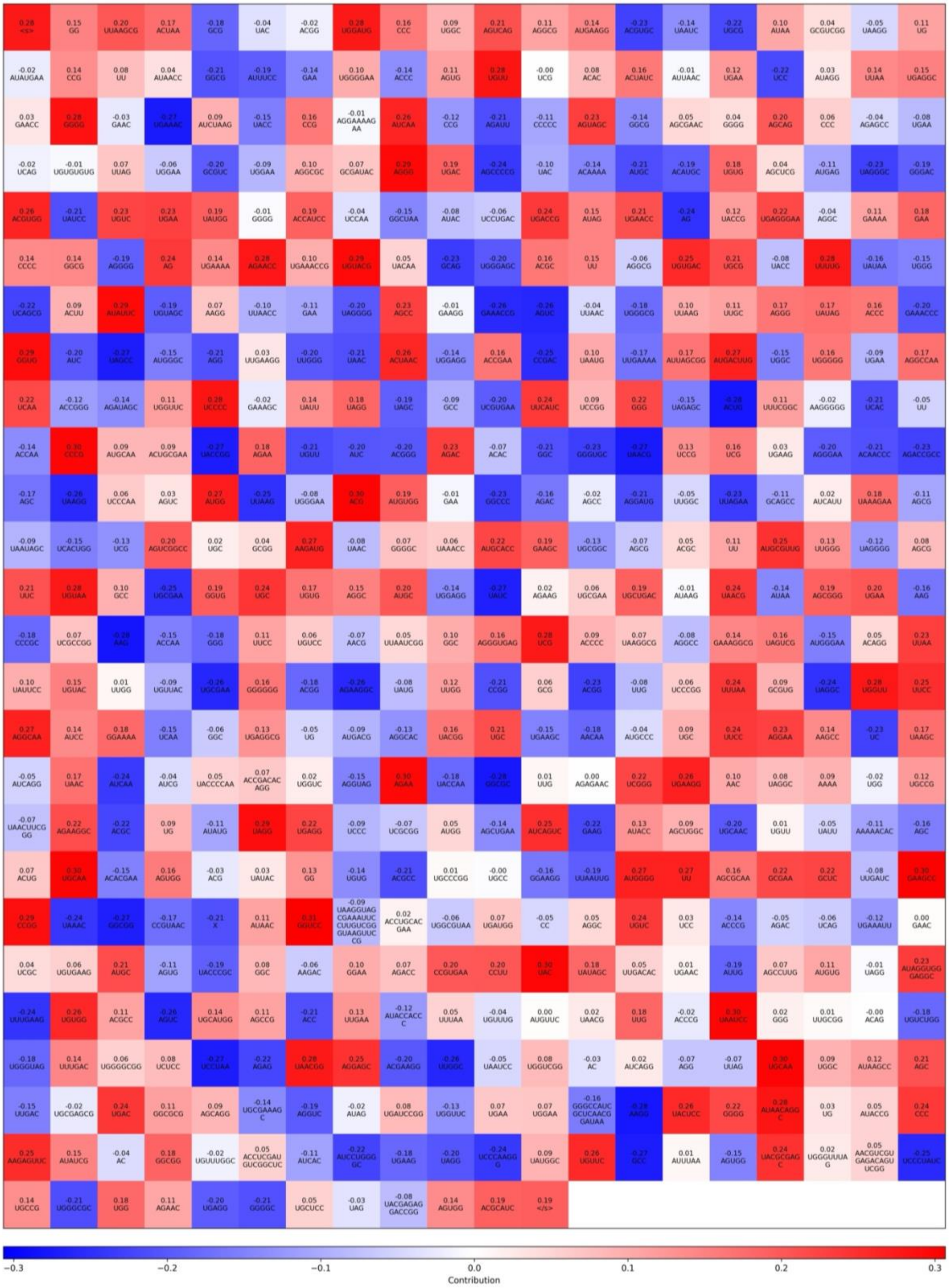


**Figure S9:** Unnormalized visualization between target “bacterial large 50S subunit ribosomal RNA” and compound “azithromycin”. In each cell, the predicted binding affinity contribution for each token and the sequence corresponding to that token can be positive, zero or negative.

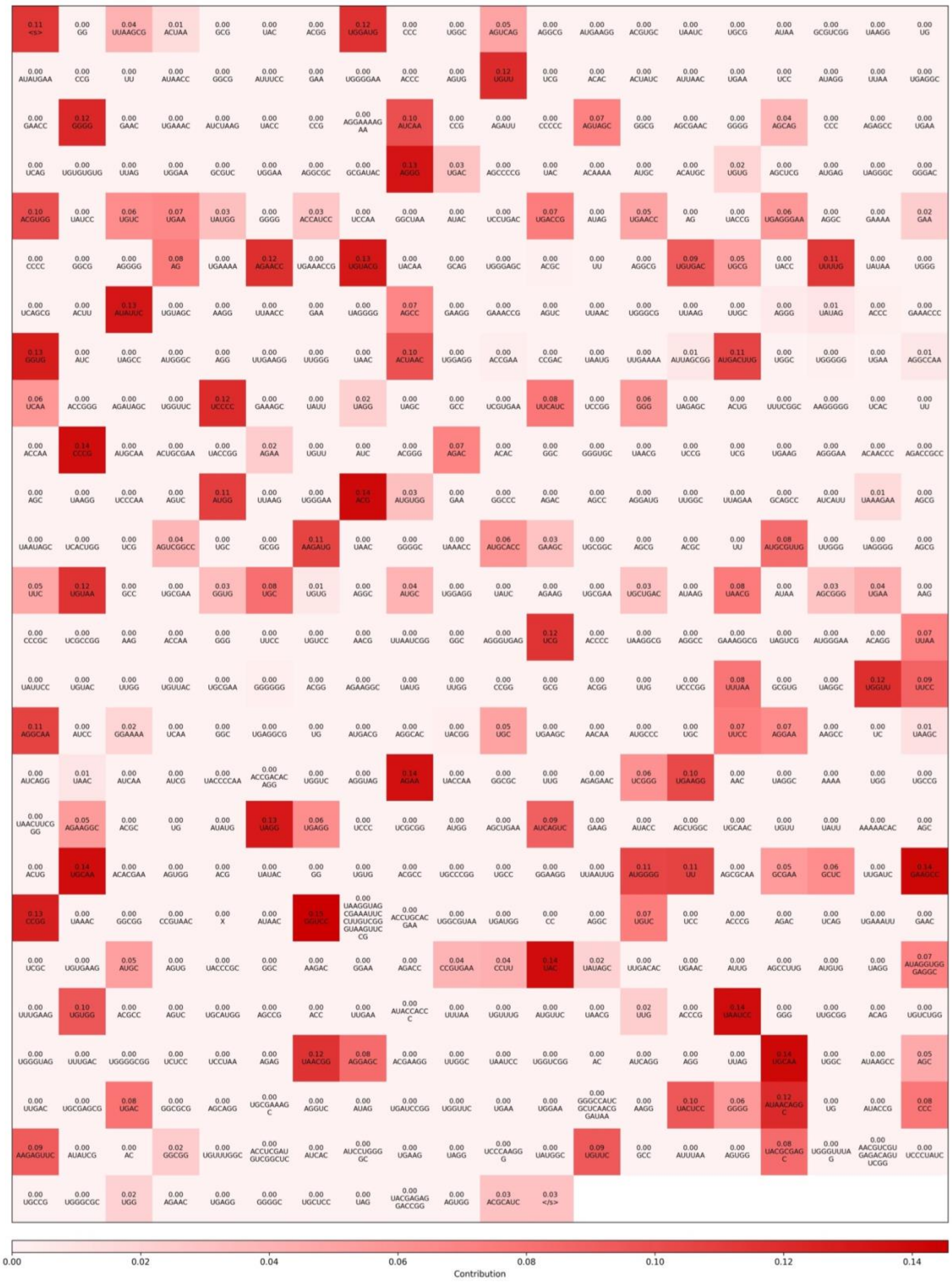

**Figure S10:** Normalized visualization between target “bacterial large 50S subunit ribosomal RNA” and compound “azithromycin”. In each cell, the predicted binding affinity contribution for each token and the sequence corresponding to that token can be only positive or zero.

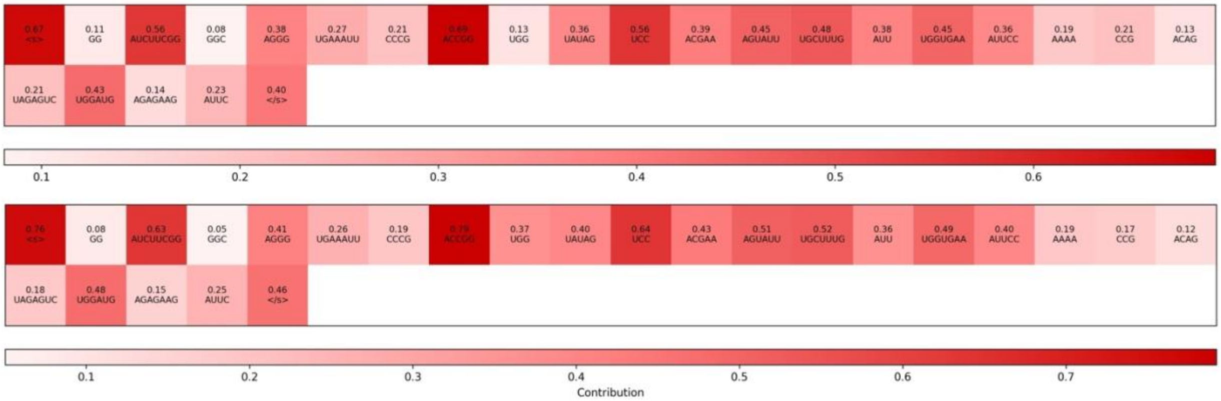


**Figure S11:** Example of normalized visualization plots showcasing the interaction contributions between tokens of the same sequence and two different compounds.

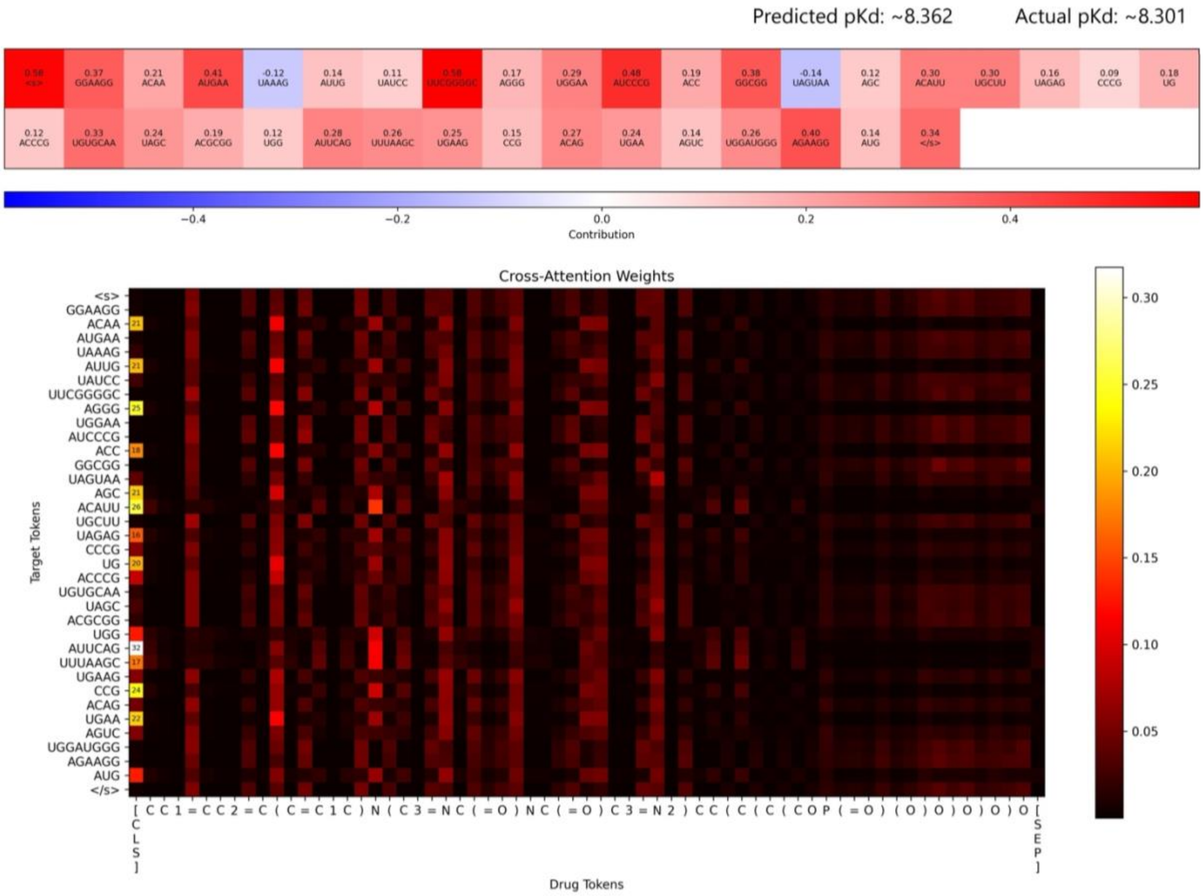


**Figure S12:** Unnormalized visualization plot and cross-attention plot between target "Bacillus subtilis's FMN riboswitch aptamer" and compound "FMN" before ablation.

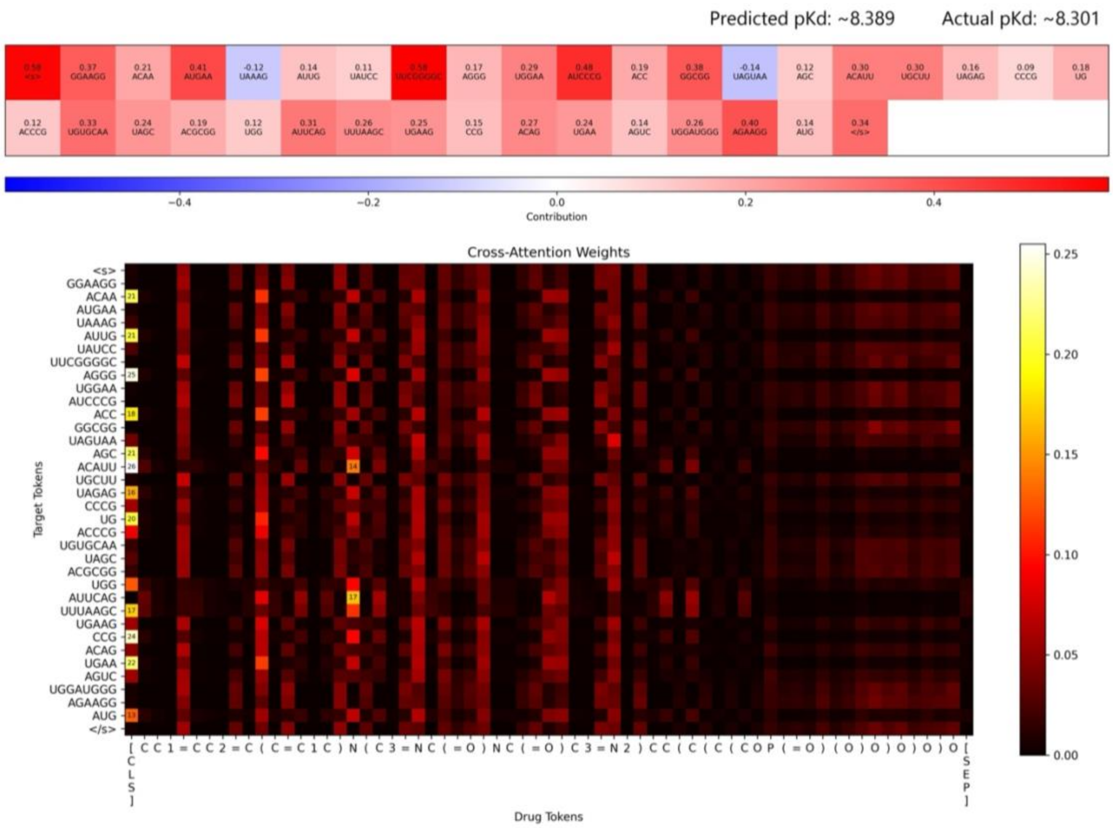


**Figure S13** Unnormalized visualization plot and cross-attention plot between target “Bacillus subtilis’s FMN riboswitch aptamer” and compound “FMN” after ablation.

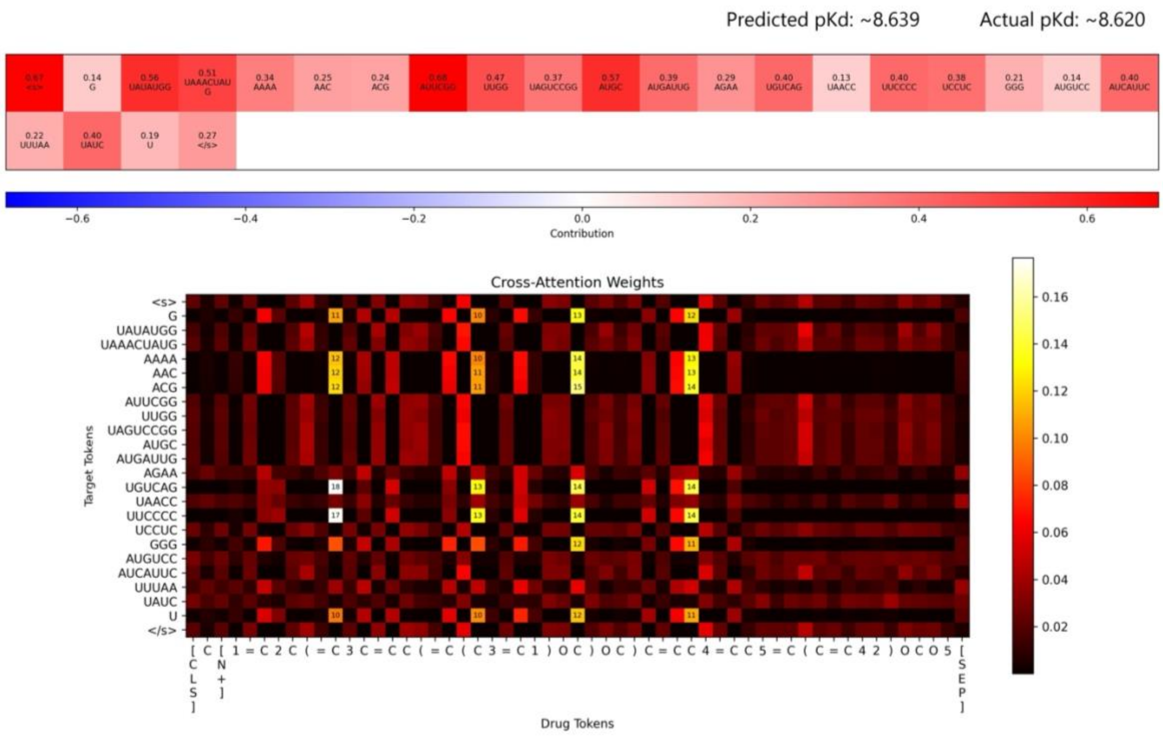

**Figure S14:** Unnormalized visualization plot and cross-attention plot between target “Bacillus subtilis’ 5’-monophosphate wt guanine riboswitch” and compound “FMN” before ablation.

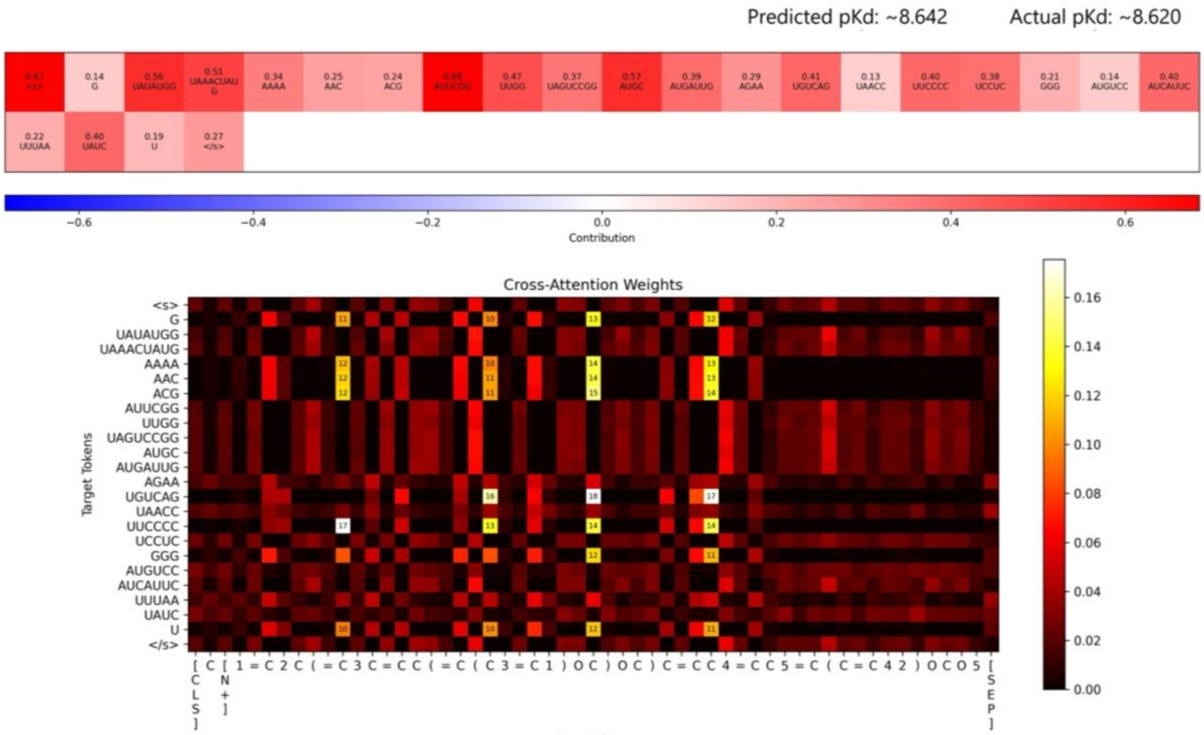


**Figure S15:** Unnormalized visualization plot and cross-attention plot between target “Bacillus subtilis’ 5’-monophosphate wt guanine riboswitch” and compound “FMN” after ablation.
